# A single cell atlas of the cycling murine ovary

**DOI:** 10.1101/2022.02.08.479522

**Authors:** ME Morris, MC Meinsohn, M Chauvin, HD Saatcioglu, A. Kashiwagi, NA. Sicher, NMP Nguyen, S Yuan, Rhian Stavely, M Hyun, PK Donahoe, B Sabatini, D Pépin

## Abstract

The estrous cycle is regulated by rhythmic endocrine interactions of the nervous and reproductive systems, which coordinate the hormonal and ovulatory functions of the ovary. Folliculogenesis and follicle progression require the orchestrated response of a variety of cell types to allow the maturation of the follicle and its sequela, ovulation, corpus luteum (CL) formation, and ovulatory wound repair. Little is known about the cell state dynamics of the ovary during the estrous cycle, and the paracrine factors that help coordinate this process. Herein we used single-cell RNA sequencing to evaluate the transcriptome of > 34,000 cells of the adult mouse ovary and describe the transcriptional changes that occur across the normal estrous cycle and other reproductive states to build a comprehensive dynamic atlas of murine ovarian cell types and states.

## Introduction

The ovary is composed of a variety of cell types that govern its dynamic functions as both an endocrine organ capable of producing hormones such as sex steroids, and a reproductive organ orchestrating the development of follicles, a structure defined by an oocyte surrounded by supporting somatic cells such as granulosa cells and theca cells. Most follicles in the ovary are quiescent primordial follicles, representing the ovarian reserve. Once activated, a primordial follicle grows in size and complexity as it progresses to primary, preantral, and antral stages, adding layers of granulosa and theca cells and forming an antral cavity, until it ultimately ejects the oocyte-cumulus complex at ovulation while the follicular remnants undergo terminal differentiation to form the corpus luteum (1). This process necessitates precise coordination of germ cells and several somatic cell types, including granulosa cells, thecal cells, vascular cells, and other stromal cells of the ovary to support the growth of the oocyte until its ovulation or follicular atresia. In addition to supporting germ cells, ovarian somatic cells must produce the necessary hormonal cues, as well as coordinate the profound tissue remodeling, necessary to accommodate these dynamic developing structures. For reproductive success to occur the state of each of these cells must change in a coordinated fashion over the course of the estrous cycle; this allows waves of follicle to grow and mature, ovulation to be triggered precisely, and provides the hormonal support necessary for pregnancy.

Single cell RNA sequencing (scRNAseq) has been used in a variety of tissues to obtain an in-depth understanding of gene expression and cellular diversity. In the ovary, this technique has allowed us, and others, to explore various physiologic processes during early ovarian development and ovarian aging (2–10). For example, Fan et al. catalogued the transcriptomic changes that occur during follicular development and regression and mapped the cell types of the human ovary using surgical specimens (9). A primate model has been used to investigate changes in cell types and states that occur in the ovary with aging (10). Zhao et al. looked at the development of the follicle during early embryonic ovarian development to discern the relationship of oocytes to their support cells in formation of follicles (2). We have used scRNAseq to identify inhibitory pathways regulated by AMH during the first wave of follicular growth in the murine ovary (8). While all these studies have helped establish a static framework to understand the major cell types in the ovary, they fail to describe the dynamic nature of cell states across the reproductive cycle, known as the estrous cycle. The estrous cycle in mice is analogous to the human menstrual cycle, which both reflect follicle development in the ovary. In mice this cycle lasts four to five days and is composed of 4 different phases known as proestrus, estrus, metestrus and diestrus. The murine proestrus is analogous to the human follicular stage and leads to ovulation at estrus. Metestrus and diestrus are analogous to early and late secretory stages of the reproductive cycle in humans, which are orchestrated by secretion of progesterone by the corpus luteum (11).

To understand more fully the dynamic effects of cyclic endocrine, autocrine, and paracrine signals on ovarian cell states we performed high-throughput single cell RNA sequencing of ovaries from adult mice across a physiological spectrum of reproductive states. Ovaries were harvested from mice in the four phases of the normal estrous cycle: proestrus, estrus, metestrus, and diestrus. Additionally, ovaries were evaluated from mice that were lactating or non-lactating 10 days postpartum mice, and from randomly cycling adult mice to increase the diversity of cell states. Herein, we describe the previously unrecognized complexity in the ovarian cellular subtypes and their cyclic expression states during the estrous cycle and identify secreted factors that represent potential marker for staging.

## Results

### scRNA-seq of adult mouse ovaries across reproductive states

To survey the dynamic transcriptional landscape of ovaries at the single cell level across a range of physiological reproductive states in sexually mature female mice, we isolated the ovaries (4 mice per group) at each stage of estrous cycling (proestrus, estrus, metestrus, diestrus), post-partum non-lactating (day 10 post-partum, with pups removed on the day they were born), post-partum lactating (day 10 postpartum, actively lactating with pups), and non-monitored mice to increase sample diversity and cell counts. Following enzymatic digestion of the ovaries, we generated single cell suspensions and sorted them by microfluidics using the inDROP methodology (15), targeting 1500 cells per animal. Resulting libraries were indexed and combined for sequencing (**Figure 1A**).

**Figure 1.**
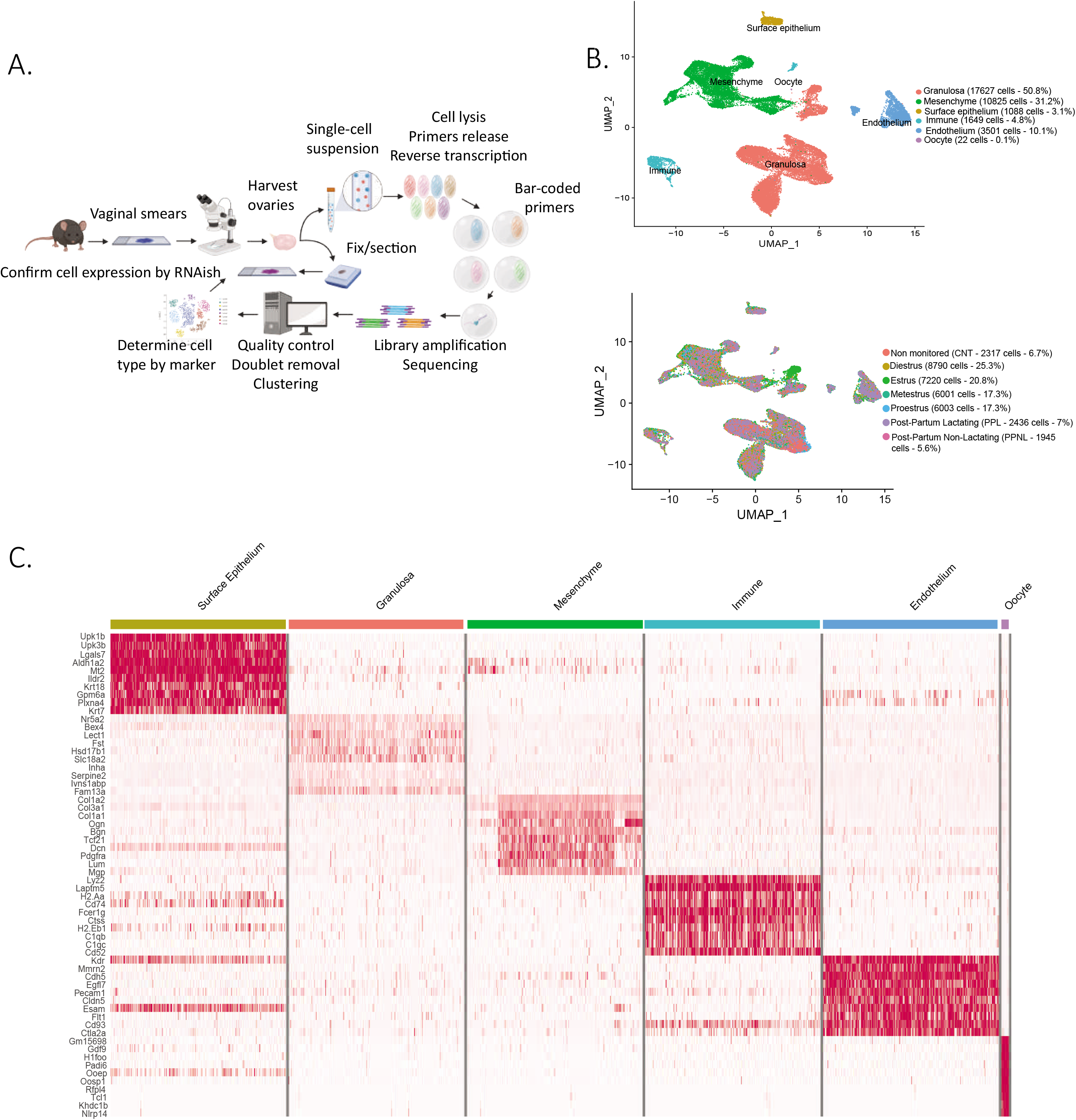
Single cell RNA sequencing of cycling mouse ovaries. A. Schematic of the single cell sequencing pipeline. B. UMAP plot featuring the different clusters of the ovary and their composition by stage of the estrus cycle, lactating stage, or unmonitored. C. Heatmap of the top 10 markers of each cluster.

Following dimensionality reduction and clustering using the Seurat algorithm we identified multiple clusters which could be combined to represent the major cell categories of the ovary (**Figure 1B**). To assign cell type identity we used cluster-specific markers which were previously described in other studies or newly identified makers later validated by RNA in situ (**Supplementary File 2**). The largest groups of clusters consisted of granulosa cells (N=17627 cells) and mesenchymal cells of the ovarian stroma (N=10825 cells). Other minor cell types were identified including endothelial cells (N= 3501 cells), ovarian surface epithelial cells (N=1088 cells), immune cells (N=1649 cells), and oocytes (N=22 cells), altogether recapitulating all the major cell types of the ovary (Figure 1 – figure supplement 1A). Oocytes were poorly represented in the dataset due to cell size limitations of inDROP, likely restricting our sampling to small oocytes of primordial follicles (**Figure 1B**). To better characterize the transcriptional signatures of the identified cell types we evaluated a heatmap of marker gene expression across the major categories of cell types and states (**Figure 1C**). Cells were also classified depending on the stage of the estrous cycle or lactating states in which the ovaries were collected (**Fig 1B**). Morphological differences between the stages of proestrus, estrus, metestrus, diestrus, and also post-partum lactating and non-lactating, were documented in **Figure 1-figure supplement 1B**. The granulosa, mesenchyme, and epithelium clusters were isolated and reanalyzed to identify subclusters.

### Single cell sequencing reveals heterogeneity within granulosa and mesenchymal cell clusters

#### Cellular diversity of Mesenchymal cells

The mesenchymal cluster was the second largest cluster identified in our analysis. Based on prior studies and conserved marker expression (9,10) we were able to identify subclusters within mesenchymal cells, and their relative abundance (percentage) as follows : early theca (16.8%), which formed the theca interna of preantral follicles, steroidogenic theca (13.2%), which formed the theca interna of antral follicles, smooth muscle cells (10.2%), which were part of the theca externa of both antral and preantral follicles, pericytes (6.2%), which surrounded the vasculature, and two interstitial stromal cell clusters, one composed of steroidogenic cells (28.7%) and the other of fibroblast-like cells (24.9%), which together constituted the bulk of the ovarian volume (outside of follicles). These subclusters can be seen in **Figure 2A**, with the top 5 expressed markers of each subcluster described in the **Figure 2B** heatmap and the top 10 listed in **Supplementary File 3.**

**Figure 2.**
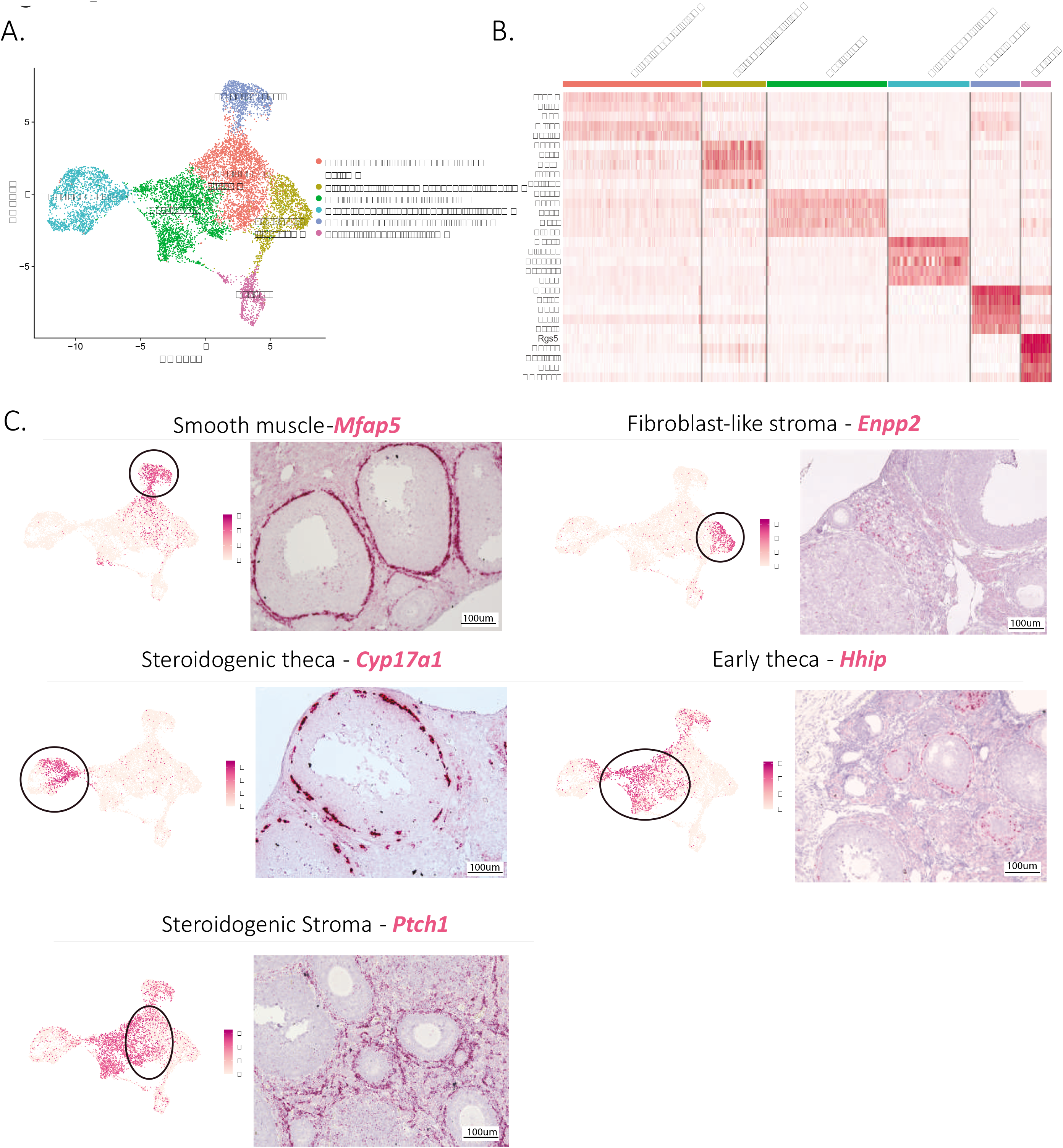
Identification of the different cell types of the mesenchyme cluster. A. UMAP plot featuring the different cell subclusters belonging to the mesenchyme cluster. B. Heatmap of the top 5 markers of each subcluster. C. Validation of the identitity of mesenchyme subcluster by UMAP-plots and RNA in situ hybridization.

Distinct transcriptional signatures were identified in each of these mesenchymal subclusters (**Figure 2B**); to confirm the presumed identity and histology of these cell types (detailed in **Figure 1-supplement 1A**), we validated markers prioritized by highest fold-change expression, highest differential percent expression, and lowest P-value (**Figure 2C**).

For the theca interna, the two clusters identified reflected the stage of development of the follicle: early thecal cells could be defined by their expression of Hedgehog Interacting Protein (*Hhip*) and were histologically associated with preantral follicles; meanwhile, the steroidogenic theca cells were identified by their expression of Cytochrome P450 Family 17 Subfamily A Member 1 (*Cyp17a1*), an essential enzyme for androgen biosynthesis (20), and were found in antral follicles (**Figure 2C**). The theca externa is a connective tissue rich in extracellular matrix situated on the outermost layer of the follicle (**Figure 1-supplement 1A**), constituted of fibroblasts, macrophages, blood vessels, and abundant smooth muscle cells which we identified based on their expression of Microfibril Associated Protein 5 (*Mfap5*) by RNA in situ hybridization (**Fig 2C**). To validate the identity and histology of these smooth muscle cells, we performed RNAish/IHC co-localization of *Mfap5* and Actin Alpha 2 (*Acta2*), another marker of smooth muscle, which confirmed their position within the theca externa. In contrast, *Hhip*, which was expressed in theca interna (both immature and steroidogenic), didn’t colocalize with Acta2 (**Figure 2-supplement 1A, B, C**). These results suggest Mfap5+ exquisitely labels smooth muscle cells of the theca externa, which are thought to perform a contractile function during ovulation (21).

Lastly, the bulk of the ovarian interstitial stromal space was made up of two closely related cell types which could not be differentiated by specific dichotomous markers, but rather distinguished based on relative expression of Ectonucleotide Pyrophosphatase/Phosphoiestrase 2 (*Enpp2*) (Fig 2C). While Enpp2+ cells represented fibroblast-like stromal cell, Enpp2-interstitial cells were enriched for expression of genes such as Patch1 (Ptch1), a member of the hedgehog signaling pathway, known as an important regulator of ovarian steroidogenesis (22), suggesting these represented steroidogenic stromal cells. Indeed, the steroidogenic activity of this stromal cell cluster was further confirmed by its high relative expression of other genes associated with steroidogenesis including Cytochrome P450 Family 11 Subfamily A Member 1 (*Cyp11a1*), Hydroxy-Delta-5-Steroid Dehydrogenase, 3 Beta- And Steroid Delta-Isomerase 1 (*Hsd3b1*), Cytochrome P450 Family 17 Subfamily A Member 1 (*Cyp17a1*), Steroid 5 Alpha-Reductase 1 (*Srd5a1*), along with other markers such as Potassium Two Pore Domain Channel Subfamily K Member 2 (*Kcnk2*) (**Figure 2-supplement 1E,F**). In contrast the fibroblast stromal cluster had enriched expression of many extracellular matrix genes such as Collagen Type I Alpha 1 Chain (*Col1a1*), Collagen Type V Alpha 1 Chain (*Col5a1*), Lumican (*Lum*), Lysyl Oxidase Like 1(*Loxl1*) (23) along with other known markers of fibroblasts such as C-X-C Motif Chemokine Ligand 14 (*Cxcl14*) (24) and WT1 Transcription Factor (*Wt1*) (25) (**Figure 2-supplement 1E,F**).

The identities of all above-described mesenchymal clusters matched previously reported markers such as Desmin (Des) for pericytes (26), Steroidogenic Acute Regulatory protein (Star) for the steroidogenic theca (27), Cellular Communication Network Factor 1 (Ccn1) for the smooth muscle (28), Receptor Activity Modifying Protein 2 (Ramp2) for the early theca cells surrounding preantral follicles (29) and finally C-X-C motif chemokine ligand 12 (Cxcl12) for both stroma clusters (30) as illustrated in **Fig2 – supplement 1D**).

#### Cellular diversity of Granulosa cells

To further explore the cellular heterogeneity within developing follicles (listed in **Figure1-supplement 1A**), we investigated the sub-clustering of granulosa cells and their transcriptional profile. Consistent with previous reports, we could distinguish discrete granulosa cell states in follicles based on their stage of development (2,9,31). Granulosa cells could be subdivided into 8 main categories: Preantral-Cumulus (27.3%), Antral-Mural (21.8%), Luteinizing mural (4.8%), Atretic (22.6%), Mitotic (14.4%), Regressing corpus luteum (CL) (3.7%), Active CL (5.4%), (**Figure 3A**). **Supplementary file 4** lists the top ten markers for each of these clusters. Distinctive gene expression programs were identified in the granulosa cell subclusters, as visualized in the heatmap (**Figure 3B**), from which we selected potential markers for validation.

**Figure 3.**
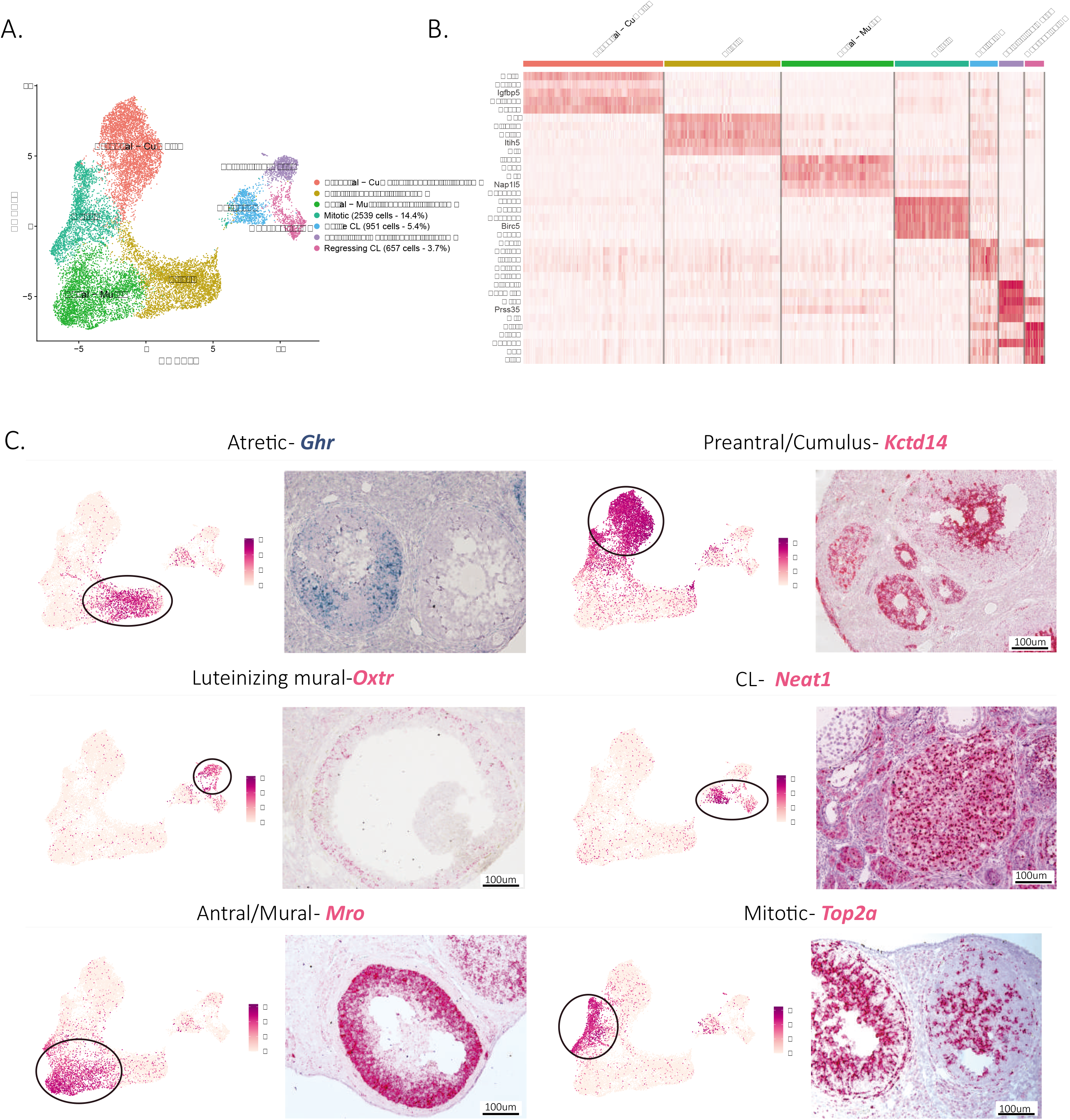
Identification of the different cell types in the granulosa cluster. A. UMAP plot featuring the different cell subclusters belonging to the granulosa cluster. B. Heatmap of the top 5 markers of each cluster. C. Validation of the identitity of granulosa subclusters by UMAP-plots RNA in situ hybridization

Early pre-antral granulosa cells, and those constituting the cumulus of antral follicles, could be identified by their shared expression of markers such as potassium channel tetramerization domain (*Kctd14*) (**Figure 3C**), which we had previously shown to be expressed by pre-antral follicles (8). In contrast, mural granulosa cells of antral follicles expressed distinct markers (**Supplementary file 4**) such as *Male-specific Transcription in the developing Reproductive Organs* (*Mro*) (**Figure 3C**). Luteinizing mural granulosa cells could be identified by the expression of previously established markers (**Supplementary File 4**) and Oxytocin Receptor gene (*Oxtr*) which we propose as a highly specific marker for this cell type, the likely target of the surge in oxytocin during estrus (32)(**Figure 3C**). Furthermore, we identified two different clusters that we hypothesize represent cell states of the corpus luteum, either active or regressing, which both expressed Nuclear Paraspeckle Assembly Transcript 1 (*Neat1*), a known marker of *CLs* (33). To confirm the active and regressing CL cell states we investigated the expression of Top2a, a mitotic marker (34), which was enriched in the active CL cluster, and Cdkn1a, a cell cycle exit and senescence marker (35), which was enriched in the regressing cluster (**Figure 3-supplement 1B, C**). Moreover, when identifying the composition of clusters depending on the reproductive stage, the regressing corpus luteum cluster was found to be composed mostly of cells derived from the post-partum non-lactating (PPNL) samples (**Figure 3-supplement 1E**), and overexpressed markers related to corpus luteum regression (36) (**Figure 3 – supplement 1F**), consistent with a post-partum anestrous state in these mice. Finally, two relatively abundant granulosa cell states could be identified based on marker expression: mitotic granulosa cells could be found in both preantral and antral follicles, and were defined by their expression of *Top2a*,and atretic granulosa cells, which expressed markers consistent with follicular atresia and apoptosis such as Phosphoinositide-3-Kinase Interacting Protein 1 (*Pik3ip1*), Nuclear Protein 1, Transcriptional Regulator (*Nupr1*), Growth Arrest And DNA Damage Inducible Alpha (*Gadd45a*), Vesicle Amine Transport 1 (*Vat1*),Transgelin (*Tagln*), Melanocyte Inducing Transcription Factor (*Mitf*) (37)(**Figure 3C, Figure 3 – supplement 1A, Supplementary File 4**). Furthermore, we propose growth hormone receptor (*Ghr*), which was highly specific to this cluster, as a specific marker of atretic follicles, which warrant further investigation of the role of growth hormone in this process (**Figure 3C**).

#### Cellular states in the ovarian surface epithelium

The epithelial cluster was composed of 1088 ovarian surface epithelium cells, which could be further subdivided into two clusters (**Figure 4A**), the larger one composed of non-dividing epithelium cells (96%) and a smaller cluster (4%) composed of mitotic epithelium characterized by proliferation markers such as Thymidine Kinase 1 (*Tk1*) (38), Rac GTPase-activating protein 1 (*Racgap1*)(39), Top2a, Casein kinase 1 (*Ck1*)(40), Protein Regulator Of Cytokinesis 1 (*Prc1*)(41), Ubiquitin Conjugating Enzyme E2 C (*Ube2c*)(42), Baculoviral IAP Repeat Containing 5 (*Birc5*)(43) (**Figure 4A, B**). Interestingly, the proliferating subcluster of ovarian surface epithelium was almost exclusively composed of cells in the estrus stage (**Figure 4C,D**), consistent with their transient amplification during ovulatory wound closure (44).

**Figure 4.**
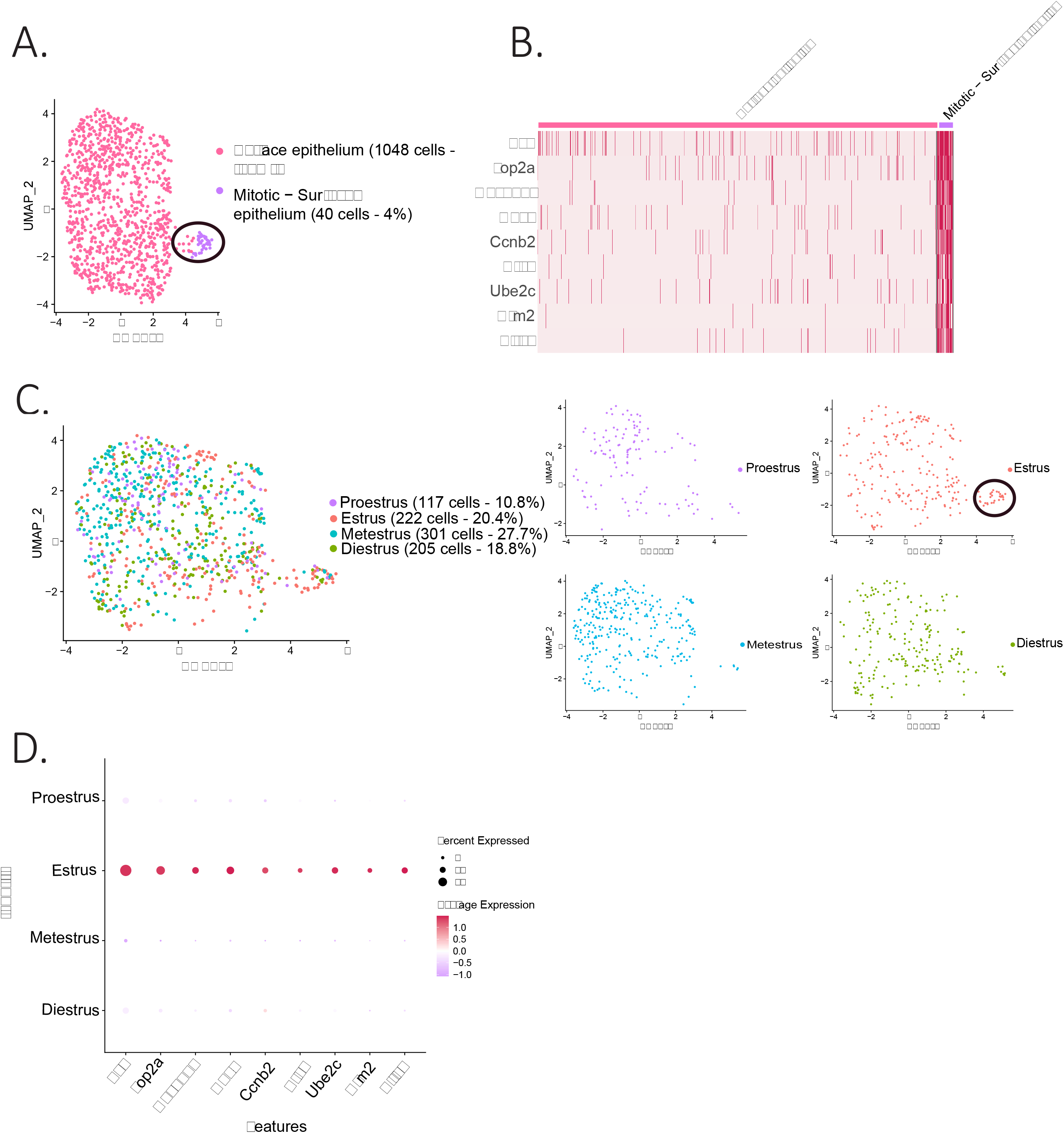
Identification of epithelial subclusters. A. UMAP plot of the epithelium cluster showing two subclusters: epithelium and proliferating epithelium. B. Heatmap of proliferation markers expressed in the proliferating epithelium cluster. C. UMAP plot of the cellular composition of the epithelium subclusters by reproductive state. D. Expression of proliferation markers depending on the phase of the estrous cycle.

### Granulosa cell transcriptome is most dynamic during the proestrus/estrus transition

To identify changes in cell states associated with the stages of the estrous cycle we focused on the granulosa cell subclusters, given the importance of follicular maturation in coordinating this process (illustrated in **Figure 1 – supplement 1B**). When comparing the composition of granulosa cell subclusters by estrous stage, we found that some clusters were dominated by cells from either the proestrous and estrous samples, particularly the clusters corresponding to “antral/mural” and “periovulatory” clusters respectively (**Figure 5A,B**). A volcano plot analysis confirmed that the transition between these two stages was characterized by 24 significantly upregulated and 10 significantly downregulated markers (**Figure 5C**), which together with the transition from estrus to metestrus, represents the largest change in gene expression. In contrast, few genes were found to significantly change during the transition from metestrus to diestrus, or diestrus to proestrus (**Figure 5 – supplement 1A**). Gene ontology analysis revealed that the most significantly differentially regulated pathways between the proestrous and estrous phases were related to ovarian matrix remodeling and steroidogenesis and hormones production (**Figure 5 – supplement 1B**). To validate the genes with significant changes in expression identified within the single cell sequencing dataset, we performed qPCR on whole-ovary samples at the proestrus to estrus transition, including the steroid biosynthesis markers Cytochrome P450 Family 19 Subfamily A Member 1 (*Cyp19a1*, p=0.0029, proestrus to estrus), Steroidogenic Acute Regulatory Protein (*Star*, p=0.0187, proestrus to estrus), Serum- and glucocorticoid-inducible kinase-1 (*Sgk1*, p=0.0056, proestrus to metestrus), as well as matrix remodeling genes such as Regulator Of Cell Cycle (*Rgcc*, p=0.0441, proestrus to estrus), Tribbles Pseudokinase 2 (*Trib2*, p=0.0023, proestrus to estrus) (**Figure 5D,E**), and immediate early genes, Fos Proto-Oncogene (*Fos*), Jun Proto-Oncogene (*Jun*, p=0.0022, proestrus to estrus), Jun Proto-Oncogene B (*Junb*, p=0.0069, proestrus to diestrus) and Early Growth Response 1 (*Egr1*, p=0.0504 estrus to diestrus), which represent a family of genes thought to be involved in wound repair, a sequela of ovulation (45–48) (**Figure 5 – supplement 1C**). Transcriptional gene expression changes were found to be concordant between the scRNAseq data and whole-ovary transcripts quantified by qPCR.

**Figure 5.**
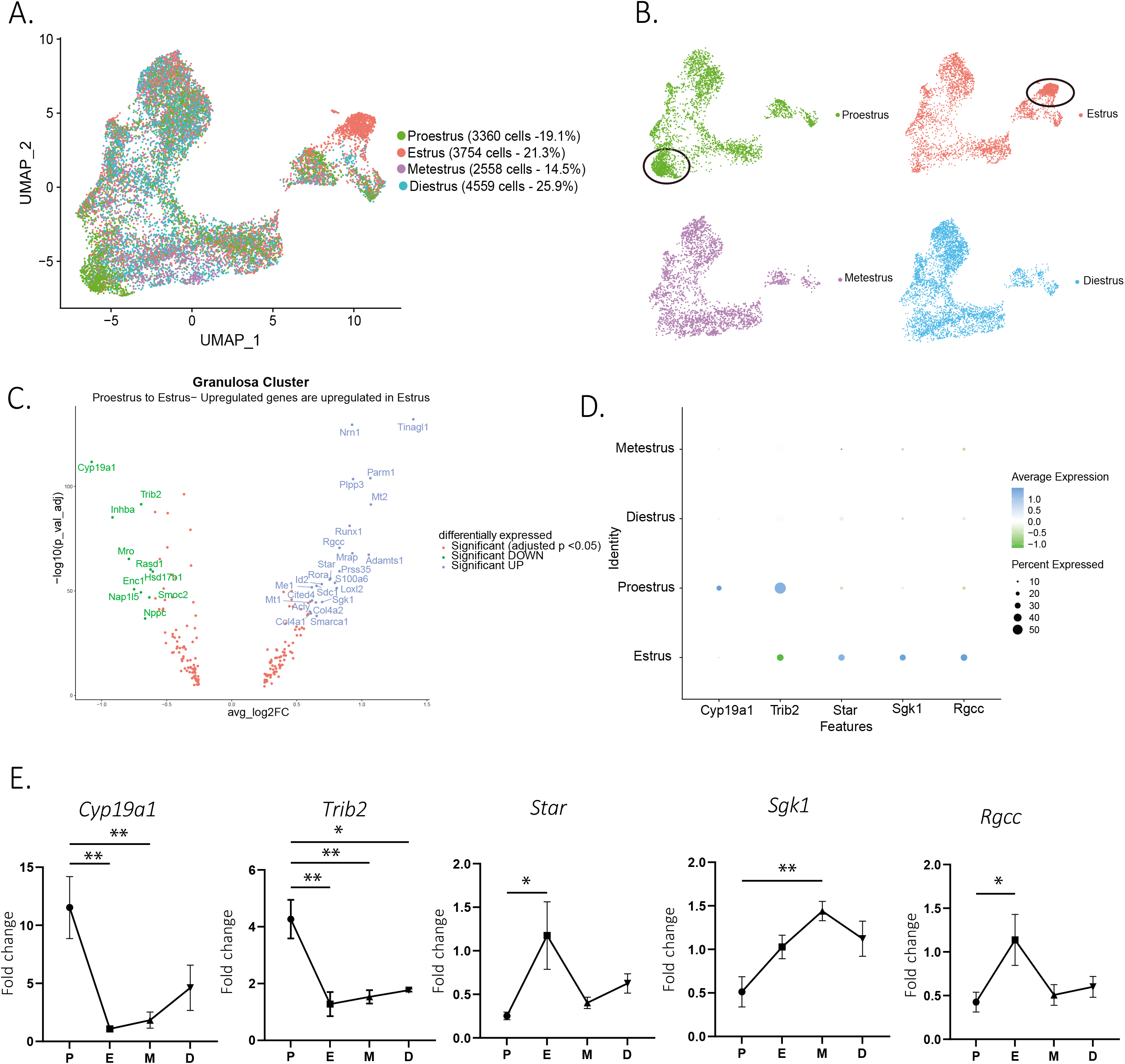
Gene expression in granulosa cells by estrous stage. A. UMAP plot featuring estrous cycle stages in the granulosa cell cluster. B. UMAP plots featuring each of the estrous cycle phases individually. C. Volcano plot of genes differentially expressed between proestrus and estrous stages. D. DotPlot of differentially expressed markers between proestrus and estrus. E. QPCR validation of differentially expressed genes involved in extracellular matrix remodeling, and steroidogenesis markers (n=5 per group, mean ± SEM, *P<0.05, **P<0.01, ***P<0.005, ****P<0.001).

### Identification and validation of secreted biomarkers varying throughout the estrous cycle

To identify new biomarkers that vary as a function of the estrous cycle and that could be used clinically for staging in reproductive medicine, we screened for differentially expressed secreted factors (DAVID Bioinformatics Resources)(49,50), which would therefore be potentially measurable in the blood. Furthermore, to ensure specificity, we prioritized genes expressed specifically in the granulosa or ovarian mesenchymal clusters and not highly expressed in other tissues based on GTEX profile (51)(**Supplementary file 5**). As a primary screen, we first validated our ability to detect gene expression changes by estrous stage using whole-ovary qPCR analysis in a sperate set of staged mice (N=4 per group). Whole ovary qPCR successfully detected expression changes of estrous cycle markers such as luteinizing hormone/choriogonadotropin receptor (52)(*Lhcgr*, p=0.0281 estrus to metestrus) and progesterone receptor (*Pgr*, p=0.0096, proestrus to estrus) (53) (**Figure 6B**). Using this method, we validated a set of significantly upregulated secreted markers in the proestrous to estrous transition, most prominent of which were natriuretic peptide C (*Nppc*, p=0.0022 proestrus to estrus) and inhibin subunit beta-A (*Inhba*, p= 0.0067, proestrus to estrus) (**Figure 6A,B**). Similarly, Tubulointerstitial Nephritis Antigen Like 1 (*Tinagl1*) and Serine Protease 35 (*Prss35*) were secreted markers significantly upregulated in estrus compared to their level of transcription in proestrus in the scRNAseq dataset (**Figure 6A**), and by qPCR (*Tinagl1* p=0.0081 proestrus to estrus, *Prss35* p=0.0008 proestrus to estrus) (**Figure 6B**). In situ RNA hybridization showed that, as expected, these markers were mostly expressed in mural granulosa cells of antral follicles, while *Nppc* was expressed in both mural and cumulus cells (**Figure 6C**).

**Figure 6.**
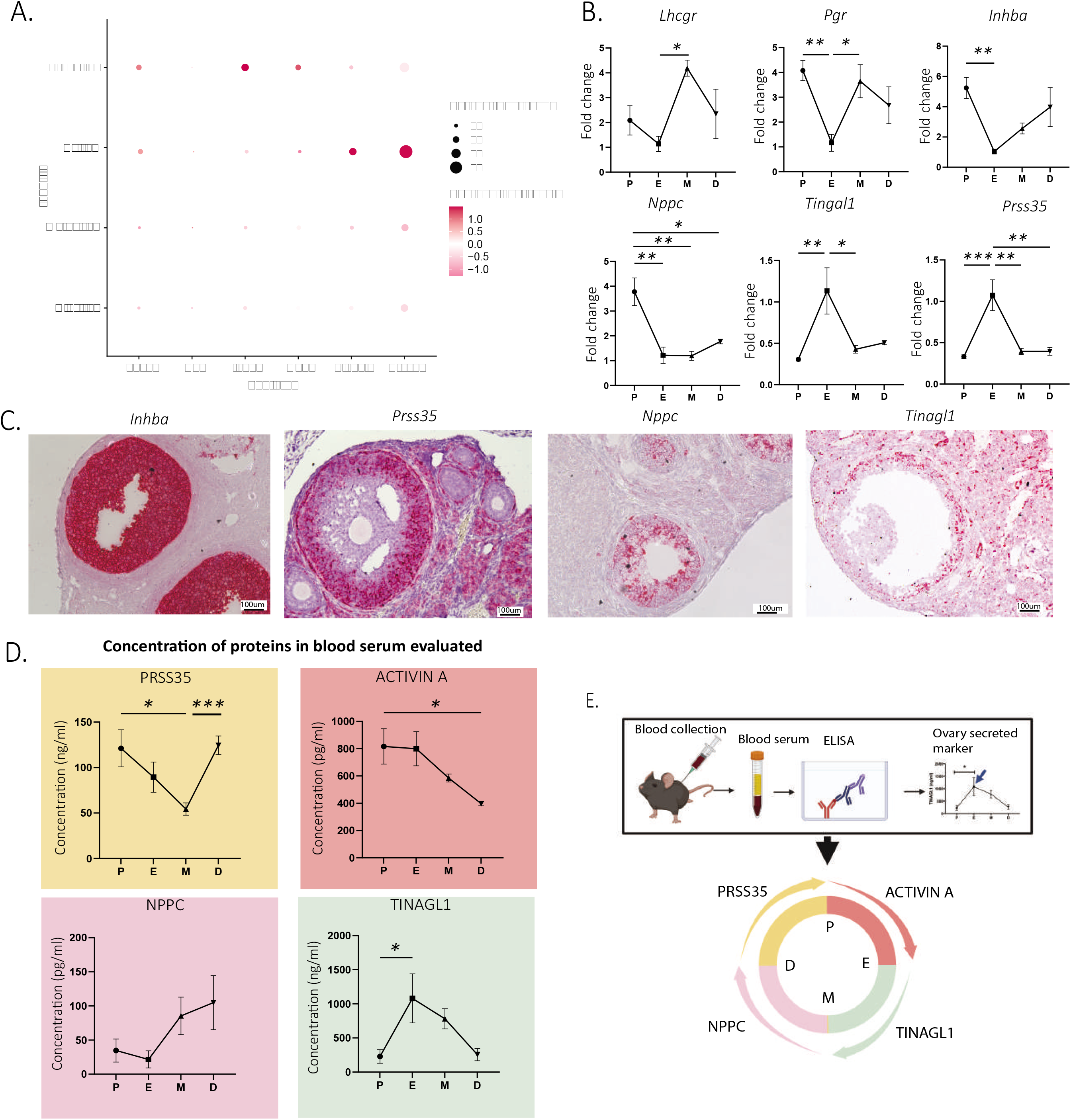
Identification and validation of new secreted estrous staging markers. A. Expression of granulosa cell transcripts varying by estrous cycle stage. B. Validation of significantly up- and downregulated transcripts of secreted estrous staging markers by qPCR (n=5 per group, mean ± SEM, *P<0.05, **P<0.01, ***P<0.005). C. Localization of estrous staging markers by in situ hybridization (RNAscope) in ovarian sections. D. Quantification of circulating estrous staging markers proteins in the blood by ELISA (n=5 per group, mean ± SEM, *P<0.05, ***P<0.005). E. Summary of the timing of expression of estrous staging markers in the blood.

To evaluate the feasibility of measuring the secreted PRSS35, NPPCC, TINAGL1 and Activin A proteins in the serum for staging we performed ELISA in mice at each stage of the estrous cycle (**Figure 6D**). We found that the Activin A concentration in the serum was significantly increased between the diestrous and proestrous stages (p=0.0312) and peaked at the proestrous stage (**Figure 6D**). The *Inhba* transcript, which encodes for the Activin and Inhibin beta-A subunit, had a similar temporal expression profile (**Figure 6B**). Circulating PRSS35 levels were lowest at the metestrous stage and were significantly increased during the transition to diestrus (p=0.0009) and remained significantly elevated until the proestrus (**Figure 6D**). In contrast, the Prss35 transcript was significantly induced earlier at estrus (**Figure 6B**). The serum concentrations of TINAGL1, which was lowest at the diestrous and proestrous stages, was significantly increased during the transition between proestrus and metestrus, peaking in estrus (p=0.0142) (**Figure 6D**). This temporal pattern of expression was recapitulated at the transcriptional level by qPCR and scRNAseq (**Figure 6A,B**). Finally, we observed a trend for serum protein concentrations of NPPC to be lowest at the proestrous and estrous stages and increase during the metestrous and diestrous stages (**Figure 6D**), although the differences were not statistically significant (p=0.0889, estrus to metestrus). Importantly, these data provide a proof of concept that four markers could be used to monitor estrous cycle progression when measured in conjunction in the blood (**Figure 6E**).

## Discussion

Single cell RNA sequencing has been used to catalog the transcriptomes of a variety of tissues in several species, across different physiological states (54). Herein, we used scRNAseq to survey the cellular diversity and the dynamic cell states of the mouse ovary across the estrous cycle and other reproductive states such as post-partum lactating and post-partum non-lactating.

The most significant changes in composition and cell states were associated granulosa cells, particularly as they cycled through the estrous stages, reflective of their important role in cyclic follicular maturation and hormone production. Early pre-antral follicle numbers are tought to be relatively stable across the estrous cycle (55), given that they are largely unresponsive to gonadotropins (56), in contrast antral follicles, whose numbers and size are more variable (55). Indeed, while subclusters such as “pre-antral granulosa cells” were equally represented in samples from proestrus, metestrus, and diestrus, others, such as the “luteinizing mural” cluster, were dominated by cells derived from one stage (in this case “estrus”). Genes enriched in this cluster had been previously reported to be involved in the ovulatory process and regulated by the LH surge, including markers of terminal differentiation and steroidogenesis such as *Smarca1* (57), *Cyp11a1* (58), Metallothionein 1 (*Mt1*), and Metallothionein 2 (*Mt2*) (59) (**Supplementary file 4**). Other genes enriched in this subcluster include Prss35 (60) and Adamts1 (61,62), which had previously been identified as playing a role in the follicular rupture necessary for ovulation.

Interestingly, we found that granulosa cells of preantral follicles and cumulus cells of antral follicles clustered together and shared markers that distinguished them from mural granulosa cells. For example, Kctd14, a member of the Potassium Channel Tetramerisation Domain Containing family, was expressed in granulosa cells during the initial growth of early pre-antral follicles, but also specifically expressed only in cumulus cells, but not mural granulosa cells, of larger antral follicles. Intriguingly, we’ve previously shown that anti-Müllerian hormone (AMH, a.k.a. MIS), which is specifically expressed by cumulus cells (63) in antral follicles, regulates KCTD14 expression in preantral follicles (8). This conservation of cellular state and marker expression from preantral granulosa cells to cumulus cells of antral follicles suggests a continuous lineage, potentially defined and maintained by the close interaction with the oocyte (63). This interpretation is consistent with the presence of a differentiation fork in the granulosa cell lineage during antrum formation, which would give rise to a distinct mural granulosa cell fate poised to respond to the LH surge. Indeed, the peri-ovulatory granulosa cell state was identified based on its expression of genes regulated by LH such as Smarca1 (57), and we propose Oxtr as a specific marker for these cells (**Figure 3-supplement 1D**). Oxtr expression was found only in the mural cells of large Graafian follicles, suggesting it indeed corresponds to an LH-stimulated mural granulosa cell state.

After ovulation, these LH-stimulated mural granulosa cells, along with the steroidogenic theca cells, terminally differentiate into luteal cells and form the corpus luteum. The corpus luteum is a transient structure with highly active steroid biosynthesis, providing the progesterone to maintain pregnancy (64). In absence of implantation the CL degenerates (65). We found this progression of the corpus luteum to be recapitulated at the transcriptional level, leading to two luteal subclusters: Active CL and Regressing CL. While the Active CL was characterized by expression of proliferation markers (Top2a) in addition to steroidogenic enzymes, the Regressing CL expressed the cell cycle inhibitor and senescence maker Cdkn1a, along with luteolysis markers such as Syndecan 4 (Sdc4), Claudin Domain Containing 1 (Cldnd1), and BTG Anti-Proliferation Factor 1 (Btg1) (36,66). The distinct expression signatures observed in these two clusters may provide insights into the molecular basis of luteolysis and warrant further investigation.

The mesenchymal cluster was also surprisingly complex and variable across the estrous cycle, reflecting its remodeling and physiological functions during follicle growth, steroid hormone production, and ovulation. For example, during follicle maturation, the ovarian stroma adjacent to the developing follicle differentiates into the theca, which is ultimately responsible for steroid hormone biosynthesis and therefore underlies the cyclic hormone production of the ovary (67). Herein we identified two thecal clusters, designated as early and steroidogenic theca. Early theca, was defined by markers such as Hhip (20,68), Mesoderm Specific Transcript (Mest) (9), and Patched 1 (Ptch1) (9,20), and given its association with small follicles, presumed to be immature and the precursor to steroidogenic theca. As the follicle matures and the antrum forms, this layer becomes more vascularized and differentiates into theca interna, which is steroidogenic. This steroidogenic theca cluster, was readily identifiable through its expression of steroidogenic enzymes such as Hydroxy-Delta-5-Steroid Dehydrogenase, 3 Beta- and Steroid Delta-Isomerase 1 (Hsd3b1), Cyp17a1, Cyp11a1, and also well-established markers such as Ferredoxin-1 (Fdx1) and prolactin receptor (Prlr) (9,69). Interestingly, we confirmed the presence of steroidogenic interstitial stromal cells which also likely contribute to sex steroid production in the ovary. Indeed, such cells likely represent the precursors of the theca interna (70,71). Smooth muscle cells, which are part of the theca externa, were identified by their expression of structural proteins such as Mfap5, Myosin Heavy Chain 11 (Myh11), Transgelin (Tagln), and Smooth Muscle Actin (Sma or Acta2) (2). In contrast to mice, human smooth muscle cells are thought to express high levels of collagen (9), which we did not observe. Another species difference between mice and humans was expression of Aldehyde Dehydrogenase 1 Family Member A1 (Aldh1a1) which we found primarily in the steroidogenic theca cluster, while it is presumably enriched in the theca externa in humans (9).

Ovulation is associated with a dramatic remodeling of the ovary, including the subsequent ovulatory wound repair. We identified fibroblast-like cells in the ovarian stroma expressing many of the extracellular matrix protein known to play a role in these processes (44,72). Another important player in ovulatory would repair is the ovarian surface epithelium (OSE), a simple mesothelial cell layer that covers the surface of the ovary, and must dynamically expand to cover wound (73,74). The OSE cluster could be identified based on well-established markers such as keratin (Krt) 7, 8 and 18 (75) and was represented by only 3% of all cells in our dataset, which could be further sub-divided in proliferative and non-proliferative states. As expected from their function in ovulatory wound repair, dividing OSE where enriched during estrus. furthermore, genes associated to wound healing such as Galectin 1 (*Lgals1*) (76), were also significantly upregulated in estrus. Similarly, the expression of the immediate-early genes Fos, Jun, Junb, and Egr1 was variable during the estrous cycle, following a common pattern of strong downregulation at estrous compared to the other stages, consistent with their temporal expression during the repair of other tissues such as the cornea (77).

Finally, to take advantage of this rich dataset, we sought to identify secreted markers which varied in abundance during the estrous cycle and could thus be used as staging biomarkers in assisted reproduction. We identified and prioritized four secreted biomarkers, expressed in mouse but also human ovaries, which varied significantly during different transitions of the estrous cycle, namely *Inhba* (78), *Prss35* (60,79), *Nppc* (80,81), and *Tinagl1* (82,83).

Activin A is a secreted protein homodimer translated from the *Inhba* transcript that is a crucial modulator of diverse ovarian functions including pituitary feedback, whose expression level depends highly on the stage of the estrous cycle (84). Quantification of Activin A protein in the serum by ELISA revealed elevated levels in the blood during both proestrus and estrus, which is consistent with studies in other species such as ewes (85). Importantly, the protein product of *Inhba*, the inhibin beta-A subunit, can be incorporated into other protein dimers, such as Activin BA, and Inhibin A, which were not measured in this study and may also represent cycling biomarkers.

The serine protease 35 transcript was expressed in the theca layers of preantral follicles and induced in granulosa cells of preovulatory follicles and all stages of the corpora lutea, peaking at the estrous stage according to qPCR, leading us to speculate that it may be involved tissue remodeling during ovulation and corpus luteum formation (60). In contrast, the PRSS35 protein levels were highest in the diestrus and proestrus stages as determined by ELISA, suggesting other tissue sources of PRSS35 or again an offset in peak protein levels due to delays in accumulation of the protein in the circulation.

The natriuretic peptide precursor C (NPPC) protein is a peptide hormone encoded by the *Npcc* gene. *Nppc* has been reported to be expressed by mural granulosa cells while its receptor Npr2 is expressed by cumulus cells(80). The pair acts on developing follicles by increasing the production of intracellular cyclic guanosine monophosphate and maintains oocyte meiotic arrest during maturation; upon downregulation of this pathway, the oocyte can escape meiotic arrest and ovulate (86). This close relationship with the ovulatory process makes *Nppc* an attractive marker to predict ovulation. Herein, qPCR analysis revealed that *Nppc* was highest in the ovary at proestrus, and was quickly and significantly downregulated at estrous, probably in response to the increased levels of LH which in turn inhibit the Nppc/Npr2 system (87). In contrast there was a trend for the circulating NPCC peptide to be highest in metestrus and diestrus, albeit not in a statistically significant way.

Finally, we evaluated the level of transcription and protein expression of the matricellular factor Tinagl1. We found both the *Tinagl1* transcript and the circulating TINAGL1 protein in the blood to be highest during estrous, thus coinciding with ovulation, with a pattern of expression consistent with expression by mural granulosa cells of antral follicles. While the role of TINAGL1 in the ovary has not been extensively investigated, it has been associated with delayed ovarian collagen deposition and increased ovulation in aging Tinagl1 knock-out mice (82).

These four potential cyclic biomarkers provide a proof of concept that a deeper understanding of changes at the single cell transcriptomics may translate into useful applications in assisted reproduction. It will be of interest to follow up the findings of cyclic expression of Activin A, PRSS35, NPPC, and TINAGL1, particularly in combination as an index, for use in humans and other species for the purpose of staging and predicting ovulation timing.

In summary, this study outlines the dynamic transcriptome of murine ovaries at the single cell level and across the estrous cycle and other reproductive states and extends our understanding of the diversity of cell types in the adult ovary. We identified herein novel biomarkers of the estrous cycle that can be readily measured in the blood and may have utility in predicting staging for assisted reproduction. This rich dataset, and extensive validation of new molecular markers of cell types of the ovary, will provide a hypothesis-generating framework of cell states across the cycle with which to elucidate the complex cellular interactions that are required for ovarian homeostasis.

## Materials and Methods

### Key Resources Table

| Key Resources Table |  |  |  |  |
| --- | --- | --- | --- | --- |
| Reagent type (species) or resource | Designation | Source or reference | Identifiers | Additional information |
| genetic reagent (*Mus musculus*) | C57BL/6-Tg(UBC-GFP)30Scha/J | Jackson Laboratory | stock #004353 |  |
| Antibody | Smooth muscle alpha action (SMA) (Rabbit polyclonal) | Abcam | #5694 | Dilution: 1:300 |
| Commercial assay or kit | ACTIVIN A commercial ELISA | RnD systems | #DAC00B |  |
| Commercial assay or kit | NPPC commercial ELISA | Novus Bio | #NBP2-75790 |  |
| Commercial assay or kit | Tinagl1 commercial ELISA | LS-Bio | #LS-F49684 |  |
| Commercial assay or kit | PRSS35 commercial ELISA | Mybiosource | #MBS9717242 |  |
| Commercial assay or kit | RNA scope 2.5 HD Duplex detection kit | ACD bio | #322500 |  |
| Commercial assay or kit | RNA scope 2.5 HD red detection kit | ACD bio | #322360 |  |
| Commercial assay or kit | the target retrieval and protease plus reagents | ACD bio | #322330 |  |
| Other | Cdkn1a (M.musculus) NM_007669.4 | ACD bio | # 408551 | RNAscope probe |
| Other | Cxcl14 (M.musculus) NM_019568.2 | ACD bio | #459741 | RNAscope probe |
| Other | Cyp11a1 (M. musculus) NM_019779.4 | ACD bio | #809181 | RNAscope probe |
| Other | Cyp17a1 (M. musculus) NM_007809.3 | ACD bio | #522611 | RNAscope probe |
| Other | Ghr (M. musculus) NM_010284.3 | ACD bio | #464951 | RNAscope probe |
| Other | Hhip (M.musculus) NM_020259.4 | ACD bio | #448441 | RNAscope probe |
| Other | Inhba (M.musculus) NM_008380.1 | ACD bio | # 455871 | RNAscope probe |
| Other | Kcnk2<br>(M.musculus)<br>NM_0011598<br>50.1 | ACD bio | #440421 | RNAscope<br>probe |
| Other | Kctd14<br>(M.musculus)<br>NM_0011362<br>35.1 | ACD bio | #517811 | RNAscope<br>probe |
| Other | Mfap5<br>(M.musculus)<br>NM_015776.2 | ACD bio | #490211 | RNAscope<br>probe |
| Other | Mro<br>(M.musculus)<br>NM_0013058<br>82.1 | ACD bio | #491181 | RNAscope<br>probe |
| Other | Neat1<br>(M.musculus)<br>NR_003513.2 | ACD bio | #440351 | RNAscope<br>probe |
| Other | Nppc<br>(M.musculus)<br>NM_010933.5 | ACD bio | # 493291 | RNAscope<br>probe |
| Other | Onecut2<br>(M.musculus)<br>NM_194268.2 | ACD bio | #520541 | RNAscope<br>probe |
| Other | Oxtr<br>(M.musculus)<br>NM_0010811<br>47.1 | ACD bio | #412171 | RNAscope<br>probe |
| Other | Prss35<br>(M.musculus)<br>NM_178738.3 | ACD bio | #492611 | RNAscope<br>probe |
| Other | Runx1 (M.<br>musculus)<br>NM_0011110<br>21.1 | ACD bio | #406671 | RNAscope<br>probe |
| Other | Tinagl1<br>(M.musculus)<br>NM_001168333 | ACD bio | #312621 | RNAscope probe |
| Other | Top2a<br>(M.musculus)<br>NM_011623.2 | ACD bio | # 491221 | RNAscope probe |
| Other | Wt1<br>(M.musculus)<br>NM_144783.2 | ACD bio | #432711 | RNAscope probe |
| Software, algorithm | R version 4.1.3 | R Project for Statistical Computing | https://scicrunch.org/resolver/SCR_001905 |  |
| Software, algorithm | Seurat package 4.1.0 | R toolkit for single cell genomics | https://satijalab.org/seurat/articles/install.html |  |
| Software, algorithm | BZ-X800 analysis software | Keyence | https://www.keyence.com/landing/microscope/lp_fluorescence.jsp |  |
| Software, algorithm | GraphPad Prism, version 9.2.0 | Graphpad |  |  |

### Mice

Animal experiments were carried out in 6-8 weeks old C57BL/6 mice obtained from Charles River Laboratory, approved by the National Institute of Health and Harvard Medical School Institutional Animal Care and Use Committee, and performed in accordance with experimental protocols 2009N000033 and 2014N000275 approved by the Massachusetts General Hospital Institutional Animal Care and Use Committee.

For the analysis of transcriptional changes in ovaries of cycling mice, animals were housed in standard conditions (12/12 hours light/dark non-inverting cycle with food and water *ad libitum*) in groups of 5 with added bedding from a cage that previously housed an adult male mouse to encourage cycling. Estrous stage was determined by observation of the vaginal opening and by vaginal swabs done at the same time daily, as previously described (12). Each mouse was monitored for a minimum of 2 weeks to ensure its cyclicity. Four mice were sacrificed in each of the 4 phases of the estrous cycle and labeled as being from experimental batch “cycling”. An additional 8 mice were included in the analysis and labeled as being from experimental batch “lactating”. Four of these mice were lactating at day 10 postpartum and four were 10 days postpartum with pups removed at delivery. Four additional mice were not monitored for cycling and included to increase sample diversity.

Additional mice were monitored throughout the estrous cycle, to collect ovaries at each stage (groups of N=5 for proestrus, estrus, metestrus, diestrus) for gene validation. Paired ovaries were collected from each staged mouse: one was used to extract mRNA for qPCR, while the other was fixed in 4% paraformaldehyde for RNAish (RNAscope™) or immunohistochemistry to validate gene expression.

### Superovulation

To stimulate superovulation, mature female mice (6-9 weeks C57BL/6) were injected IP with 5 IU of pregnant mare serum gonadotropin (PMSG) (Calbiochem, San Diego, CA), followed 48 hours later by 5 IU of human chorionic gonadotropin (hCG; Millipore Sigma, St. Louis, MO). The mice were euthanized 8 hours after hCG treatment and ovaries harvested.

### Staging of estrous cycle by vaginal cytology

As previously described (12,13), staging of mice was performed using a wet cotton swab, introduced into the vaginal orifice then smeared onto a glass slide which was air-dried, stained with Giemsa, and scored for cytology by two independent observers. Briefly, proestrus was determined if the smear showed a preponderance of nucleated epithelial cells as well as leukocytes. Estrous was marked by an abundance of cornified epithelial cells, while metestrous smears contained a mixture of cornified epithelial cells and leukocytes. Finally, diestrus was characterized by abundant leukocytes with low numbers of cornified epithelium or nucleated epithelial cells.

### Generation of single cell suspension

Single cell suspension from mouse ovaries were obtained as previously described with uterine enzymatic dissociation (14). Briefly, ovaries were incubated for 30min at 34C in dissociation medium (82 mM Na_2_SO_4_, 30 mM K_2_SO_4_, 10 mM Glucose, 10 mM HEPES, and 5 mM MgCL_2_, pH 7.4) containing 15 mg of Protease XXIII (Worthington), 100 U Papain, with 5 mm L-Cysteine, 2.5 mM EDTA (Worthington) and 1333 U of DNase 1 (Worthington). The reaction was then stopped in cold medium, and samples were mechanically dissociated, filtrated, and spun down three times before being resuspended to a concentration of 150,000 cells/mL in 20% Optiprep (Sigma) for inDrop sorting.

### Single-cell RNA sequencing (inDrops)

Fluidic sorting was performed using the inDrop platform at the Single Cell Core facility at Harvard Medical School as previously described (15,16). We generated libraries of approximately 1500 cells per animal which were sequenced using the NextSeq500 (Illumina) platform. Transcripts were processed according to a previously published pipeline (15) used to build a custom transcriptome from the Ensemble GRCm38 genome and GRCm38.84 annotation using Bowtie 1.1.1. Unique molecular identifiers (UMIs) were used to reference sequence reads back to individual captured molecules, referred to as UMIFM counts. All steps of the pipeline were run using default parameters unless explicitly specified.

### Single-cell RNAseq data analysis

#### Data processing

The initial Seurat object was created using thresholds to identify putative cells (unique cell barcodes) with the following parameters: 1000-20000 unique molecular identifiers, 500-5000 genes, and less than 15% mitochondrial genes. The final merged dataset contained ~70000 cells which were clustered based on expression of marker genes. These were further processed in several ways to exclude low-quality data and potential doublets. Visualization of single-cell data was performed using a nonlinear dimensionality-reduction technique, uniform manifold approximation and projection (UMAP). Markers for each level of cluster were identified using MAST in Seurat (R version 4.1.3 - Seurat version 4.1.0). Following identification of the main clusters (granulosa, mesenchyme, endothelium, immune, epithelium, and oocyte) we reanalyzed each cluster population to perform subclustering. Briefly, the granulosa, mesenchyme, and epithelium clusters were extracted from the integrated dataset by the Subset function. The isolated cluster was then divided into several subclusters following normalization, scale, principal component analysis, and dimensionality reductions as previously described (17).

#### Volcano Plots

Highly differentially expressed genes between different estrous cycles were identified using the function FindMarkers in Seurat. Volcano plots were generated using ggplot2 package in R.

#### Pathway Enrichment Analysis

Differentially expressed genes with at least two-fold changes between contiguous estrous stages were used as input for Gene Ontology Enrichment Analysis by clusterProfiler. Enrichplot package was used for visualization. Biological process sub-ontology was chosen for this analysis.

#### Principal component analysis (PCA)

PCA was used to identify common patterns of gene expression across stages of the cycle. For each Level0 cluster object, cycling cells were extracted and genes that were expressed in more than 5% of cells were identified. The expression of these genes in the cycling cells were scaled (set to mean zero, standard deviation 1) and averaged across each of the 4 cycle stages. PCA was run (prcomp) on the average scaled expression data.

#### Data availability

The scRNAseq count matrix dataset was deposited in OSF (18), and at the Broad Institute Single Cell Portal under study number SCP1914 (19).

### In situ hybridization and immunohistochemistry

In situ hybridizations were performed using ACDBio kits as per manufacturer’s protocol, and as previously described (14) Briefly, RNAish was developed using the RNAscope® 2.5 HD Reagent Kit (RED and Duplex, ACD Bio). Following deparaffinization in xylene, dehydration, peroxidase blocking, and heat-induced epitope retrieval by the target retrieval and protease plus reagents (ACD bio), tissue sections were hybridized with probes for the target genes (see Key resources material table for accession number and catalog number of each gene) in the HybEZ hybridization oven (ACD Bio) for 2 hours at 40°C. The slides were then processed for standard signal amplification steps, and chromogen development. Slides were counterstained in 50% hematoxylin (Dako), air dried, and cover-slipped with EcoMount™. In addition to cycling and noncycling mice, superovulated mice were used to validate markers from follicles associated with LH surge response in ovulatory follicles at the estrous stage for more precise timing of collection.

For colocalization of RNAish staining with immunohistochemistry, we first processed the tissue section for RNAscope™ as described above, including deparaffinization, antigen retrieval, hybridization, and chromogen development. Sections were then blocked in 3% bovine serum albumin (BSA) in Tris-buffered solution (TBS) for 1 hr. Following 3 washes with TBS, the sections were incubated with the primary antibody (smooth muscle actin primary antibody (SMA); 1:300, Abcam) overnight at 4°C and developed with Dako EnVision + System horseradish peroxidase (HRP). Labeled Polymer Anti-Rabbit was used as the secondary antibody, and the HRP signal was detected using the Dako detection system. Slides were then counterstained in hematoxylin and mounted as described above.

### RT-qPCR

Mice were monitored through the estrous cycle and sacrificed at specific stage/time-points as described above. Ovaries were dissected and total RNA was extracted using the Qiagen RNA extraction kit (Qiagen). A cDNA library was synthesized from 500 ng total RNA using SuperScript III First-Strand Synthesis System for RT-PCR using manufacturer’s instructions with random hexamers (Invitrogen). The primers used for this study are described in **Supplementary File 1**. Expression levels were normalized to the Gapdh transcript using cycle threshold (Ct) values logarithmically transformed by the 2-ΔCt function.

### ELISA

Blood was collected from mice by facial vein puncture, incubated at RT until spontaneously clotted, centrifuged at 8000rpm for 5min to collect the serum layer, and diluted 1/10 in each ELISA kit according to the manufacturing protocol; Mouse CNP/NPPC ELISA kit; Mouse serine protease inactive 35 (PRSS35) ELISA kit; Mouse TINAGL1 / Lipocalin 7 ELISA kit; and Human/Mouse/Rat Activin A Quantikine ELISA Kit (see Key resources material table).

---

**Figure 1 – supplement 1 – Ovarian morphology by reproductive state**. A. Illustration of the different cell types of the ovary. B. Representative micrographs of sections of ovaries at each stage of the estrous cycle, post-partum lactating and post-partum non-lactating reproductive states.

**Figure 2 – supplement 1 – Characterization of mesenchymal cell clusters**. A. Co-expression of Acta2 and Mfap5 or Acta2 and Hhip in UMAP plots. Co-localization of B. Acta2 and Mfap5 or C. Acta2 and Hhip in ovarian tissue sections stained by IHC and RNA in situ hybridization. D. Expression of mesenchymal markers by cluster in DotPlot. E. Expression of steroidogenesic and fibroblast markers in DotPlot that differ between the two interstitial stromal cell clusters. F. RNA in situ hybridization of different markers representative of the steroidogenic stroma (Cyp11a1, Ptch1) and the fibroblast-like stroma cell clusters (Kcnk2, Cxcl14, and Col1a1).

**Figure 3-supplement 1 – Characterization of apoptotic and corpus luteum clusters.** A. UMAP plots of different apoptotic markers that colocalize with *Ghr* in the apoptotic cluster. B. Colocalization of Top2a and Cdkn1a in UMAP plots of corpus luteum clusters. C. RNA in situ hybridization of Top2a and Cdkn1a in ovarian sections. Representative active (Top2a+) and regressing (Cdkn1a+) corpora lutea. D. Examples of luteinizing mural markers in DotPlot that differ from antral mural. E. UMAP plot of cellular distribution of post-partum lactating (PPL) cells across the active corpus luteum cluster and post-partum non-lactating (PPNL) cells across the regressing corpus luteum cluster. F. Heatmap of representative markers of active and regressing corpora lutea.

**Figure 5 – supplement 1 – Characterization of the granulosa cell transcriptome across the estrous cycle**. A. Volcano plot of genes differentially expressed between the estrous/metestrous, metestrous/diestrous and diestrous/proestrous stages. B. Pathway analysis of differentially expressed genes (DEG) between the different stages of the estrous cycle. The x-axis shows the gene ratio (the percentage of total DEGs in the given GO terms). “Count”, reflected by the dot size, represents the number of genes enriched in a GO term and dot color represents the adjusted p values. . C. QPCR validation of ovarian surface epithelium-expressed markers (n=5 per group, mean ± SEM, *P<0.05, **P<0.01, ****P<0.001).

**Supplementary File 1 - List of primers used for qPCR experiments**

**Supplementary File 2 – Top 10 markers expressed in each ovary cluster**

**Supplementary File 3 - Top 10 markers from each mesenchyme subclusters**

**Supplementary File 4 - Top 10 markers from each granulosa subclusters**

**Supplementary File 5 - Secreted markers expressed in granulosa cells varying with the estrous cycle**

**Source data 1 – qPCR experiments individual p values**

**Source data 2 – ELISA experiments individual p values**

## Acknowledgements

We thank LiHua Zhang, Bugra Uluyurt, Phoebe A. May, Caroline Coletti, and Sarah Mustafa Eisa for technical help. This study was supported by the National Institute for Child Health and Human Development to D.P. (1R01HD102014-01), the Huiying Fellowship (H.D.S.), Sudna Gar Fellowship (D.P.), Massachusetts General Hospital Executive Committee on Research (D.P. and P.K.D.), and royalties (P.K.D.) from the use of the MIS ELISA in infertility clinics.

## Bibliography

1. Dunlop CE, Anderson RA. The regulation and assessment of follicular growth. Scand J Clin Lab Invest Suppl 2014;244:13–17; discussion 17.

2. Zhao Z-H, Ma J-Y, Meng T-G, Wang Z-B, Yue W, Zhou Q, Li S, Feng X, Hou Y, Schatten H, Ou X-H, Sun Q-Y. Single-cell RNA sequencing reveals the landscape of early female germ cell development. FASEB J 2020;34(9):12634–12645.

3. Stévant I, Kühne F, Greenfield A, Chaboissier M-C, Dermitzakis ET, Nef S. Dissecting Cell Lineage Specification and Sex Fate Determination in Gonadal Somatic Cells Using Single-Cell Transcriptomics. Cell Rep 2019;26(12):3272–3283.e3.

4. Wagner M, Yoshihara M, Douagi I, Damdimopoulos A, Panula S, Petropoulos S, Lu H, Pettersson K, Palm K, Katayama S, Hovatta O, Kere J, Lanner F, Damdimopoulou P. Single-cell analysis of human ovarian cortex identifies distinct cell populations but no oogonial stem cells. Nat Commun 2020;11(1):1147.

5. Niu W, Spradling AC. Two distinct pathways of pregranulosa cell differentiation support follicle formation in the mouse ovary. Proc Natl Acad Sci U S A 2020;117(33):20015–20026.

6. Jevitt A, Chatterjee D, Xie G, Wang X-F, Otwell T, Huang Y-C, Deng W-M. A single-cell atlas of adult Drosophila ovary identifies transcriptional programs and somatic cell lineage regulating oogenesis. PLoS Biol 2020;18(4):e3000538.

7. Man L, Lustgarten-Guahmich N, Kallinos E, Redhead-Laconte Z, Liu S, Schattman B, Redmond D, Hancock K, Zaninovic N, Schattman G, Rosenwaks Z, James D. Comparison of Human Antral Follicles of Xenograft versus Ovarian Origin Reveals Disparate Molecular Signatures. Cell Rep 2020;32(6):108027.

8. Meinsohn M-C, Saatcioglu HD, Wei L, Li Y, Horn H, Chauvin M, Kano M, Nguyen NMP, Nagykery N, Kashiwagi A, Samore WR, Wang D, Oliva E, Gao G, Morris ME, Donahoe PK, Pépin D. Single-cell sequencing reveals suppressive transcriptional programs regulated by MIS/AMH in neonatal ovaries. Proc Natl Acad Sci U S A 2021;118(20). doi:10.1073/pnas.2100920118.

9. Fan X, Bialecka M, Moustakas I, Lam E, Torrens-Juaneda V, Borggreven NV, Trouw L, Louwe LA, Pilgram GSK, Mei H, van der Westerlaken L, Chuva de Sousa Lopes SM. Single-cell reconstruction of follicular remodeling in the human adult ovary. Nat Commun 2019;10(1):3164.

10. Wang S, Zheng Y, Li J, Yu Y, Zhang W, Song M, Liu Z, Min Z, Hu H, Jing Y, He X, Sun L, Ma L, Esteban CR, Chan P, Qiao J, Zhou Q, Izpisua Belmonte JC, Qu J, Tang F, Liu G-H. Single-Cell Transcriptomic Atlas of Primate Ovarian Aging. Cell 2020;180(3):585–600.e19.

11. Ajayi AF, Akhigbe RE. Staging of the estrous cycle and induction of estrus in experimental rodents: an update. Fertil Res Pract 2020;6:5.

12. Kano M, Sosulski AE, Zhang L, Saatcioglu HD, Wang D, Nagykery N, Sabatini ME, Gao G, Donahoe PK, Pépin D. AMH/MIS as a contraceptive that protects the ovarian reserve during chemotherapy. Proc Natl Acad Sci U S A 2017;114(9):E1688–E1697.

13. Byers SL, Wiles MV, Dunn SL, Taft RA. Mouse estrous cycle identification tool and images. PLoS One 2012;7(4):e35538.

14. Saatcioglu HD, Kano M, Horn H, Zhang L, Samore W, Nagykery N, Meinsohn M-C, Hyun M, Suliman R, Poulo J, Hsu J, Sacha C, Wang D, Gao G, Lage K, Oliva E, Morris Sabatini ME, Donahoe PK, Pépin D. Single-cell sequencing of neonatal uterus reveals an Misr2+endometrial progenitor indispensable for fertility. Elife 2019;8. doi:10.7554/eLife.46349.

15. Klein AM, Mazutis L, Akartuna I, Tallapragada N, Veres A, Li V, Peshkin L, Weitz DA, Kirschner MW. Droplet barcoding for single-cell transcriptomics applied to embryonic stem cells. Cell 2015;161(5):1187–1201.

16. Macosko EZ, Basu A, Satija R, Nemesh J, Shekhar K, Goldman M, Tirosh I, Bialas AR, Kamitaki N, Martersteck EM, Trombetta JJ, Weitz DA, Sanes JR, Shalek AK, Regev A, McCarroll SA. Highly Parallel Genome-wide Expression Profiling of Individual Cells Using Nanoliter Droplets. Cell 2015;161(5):1202–1214.

17. Niu W, Spradling AC. Two distinct pathways of pregranulosa cell differentiation support follicle formation in the mouse ovary. Proc. Natl. Acad. Sci. U.S.A. 2020. doi:10.1073/pnas.2005570117.

18. Pepin. “Single Cell Sequencing of the Mouse Ovary in Diverse Reproductive States.” OSF. 2022.

19. Single Cell Portal. Available at: https://singlecell.broadinstitute.org/single_cell/study/SCP1914/a-single-cell-atlas-of-the-cycling-murine-ovary. Accessed July 7, 2022.

20. Richards JS, Ren YA, Candelaria N, Adams JE, Rajkovic A. Ovarian Follicular Theca Cell Recruitment, Differentiation, and Impact on Fertility: 2017 Update. Endocr Rev 2018;39(1):1–20.

21. Young JM, McNeilly AS. Theca: the forgotten cell of the ovarian follicle. Reproduction 2010;140(4):489–504.

22. Spicer LJ, Sudo S, Aad PY, Wang LS, Chun S-Y, Ben-Shlomo I, Klein C, Hsueh AJW. The hedgehog-patched signaling pathway and function in the mammalian ovary: a novel role for hedgehog proteins in stimulating proliferation and steroidogenesis of theca cells. Reproduction 2009;138(2):329–339.

23. Muhl L, Genové G, Leptidis S, Liu J, He L, Mocci G, Sun Y, Gustafsson S, Buyandelger B, Chivukula IV, Segerstolpe Å, Raschperger E, Hansson EM, Björkegren JLM, Peng X-R, Vanlandewijck M, Lendahl U, Betsholtz C. Single-cell analysis uncovers fibroblast heterogeneity and criteria for fibroblast and mural cell identification and discrimination. Nat Commun 2020;11(1):3953.

24. Lu J, Chatterjee M, Schmid H, Beck S, Gawaz M. CXCL14 as an emerging immune and inflammatory modulator. J Inflamm (Lond) 2016;13:1.

25. He H, Luthringer DJ, Hui P, Lau SK, Weiss LM, Chu PG. Expression of CD56 and WT1 in ovarian stroma and ovarian stromal tumors. Am J Surg Pathol 2008;32(6):884–890.

26. Hughes S, Chan-Ling T. Characterization of smooth muscle cell and pericyte differentiation in the rat retina in vivo. Invest Ophthalmol Vis Sci 2004;45(8):2795–2806.

27. Kiriakidou M, McAllister JM, Sugawara T, Strauss JF 3rd. Expression of steroidogenic acute regulatory protein (StAR) in the human ovary. J Clin Endocrinol Metab 1996;81(11):4122–4128.

28. Yang R, Chen Y, Chen D. Biological functions and role of CCN1/Cyr61 in embryogenesis and tumorigenesis in the female reproductive system (Review). Mol Med Rep 2018;17(1):3–10.

29. Hatzirodos N, Hummitzsch K, Irving-Rodgers HF, Rodgers RJ. Transcriptome comparisons identify new cell markers for theca interna and granulosa cells from small and large antral ovarian follicles. PLoS One 2015;10(3):e0119800.

30. Porcile C, Bajetto A, Barbieri F, Barbero S, Bonavia R, Biglieri M, Pirani P, Florio T, Schettini G. Stromal cell-derived factor-1alpha (SDF-1alpha/CXCL12) stimulates ovarian cancer cell growth through the EGF receptor transactivation. Exp Cell Res 2005;308(2):241–253.

31. Gallardo TD, John GB, Shirley L, Contreras CM, Akbay EA, Haynie JM, Ward SE, Shidler MJ, Castrillon DH. Genomewide discovery and classification of candidate ovarian fertility genes in the mouse. Genetics 2007;177(1):179–194.

32. Ho ML, Lee JN. Ovarian and circulating levels of oxytocin and arginine vasopressin during the estrous cycle in the rat. Acta Endocrinol (Copenh) 1992;126(6):530–534.

33. Nakagawa S, Shimada M, Yanaka K, Mito M, Arai T, Takahashi E, Fujita Y, Fujimori T, Standaert L, Marine J-C, Hirose T. The lncRNA Neat1 is required for corpus luteum formation and the establishment of pregnancy in a subpopulation of mice. Development 2014;141(23):4618–4627.

34. Donadeu FX, Fahiminiya S, Esteves CL, Nadaf J, Miedzinska K, McNeilly AS, Waddington D, Gérard N. Transcriptome profiling of granulosa and theca cells during dominant follicle development in the horse. Biol Reprod 2014;91(5):111.

35. Ock S-A, Knott JG, Choi I. Involvement of CDKN1A (p21) in cellular senescence in response to heat and irradiation stress during preimplantation development. Cell Stress Chaperones 2020;25(3):503–508.

36. Talbott H, Hou X, Qiu F, Zhang P, Guda C, Yu F, Cushman RA, Wood JR, Wang C, Cupp AS, Davis JS. Early transcriptome responses of the bovine midcycle corpus luteum to prostaglandin F2α includes cytokine signaling. Mol Cell Endocrinol 2017;452:93–109.

37. Terenina E, Fabre S, Bonnet A, Monniaux D, Robert-Granié C, SanCristobal M, Sarry J, Vignoles F, Gondret F, Monget P, Tosser-Klopp G. Differentially expressed genes and gene networks involved in pig ovarian follicular atresia. Physiol Genomics 2017;49(2):67–80.

38. Liu C, Wang J, Zhao L, He H, Zhao P, Peng Z, Liu F, Chen J, Wu W, Wang G, Dong F. Knockdown of Thymidine Kinase 1 Suppresses Cell Proliferation, Invasion, Migration, and Epithelial-Mesenchymal Transition in Thyroid Carcinoma Cells. Front Oncol 2019;9:1475.

39. Yang X-M, Cao X-Y, He P, Li J, Feng M-X, Zhang Y-L, Zhang X-L, Wang Y-H, Yang Q, Zhu L, Nie H-Z, Jiang S-H, Tian G-A, Zhang X-X, Liu Q, Ji J, Zhu X, Xia Q, Zhang Z-G. Overexpression of Rac GTPase Activating Protein 1 Contributes to Proliferation of Cancer Cells by Reducing Hippo Signaling to Promote Cytokinesis. Gastroenterology 2018;155(4):1233–1249.e22.

40. Gao H-S, Lin S-Y, Han X, Xu H-Z, Gao Y-L, Qin Z-Y. Casein kinase 1 (CK1) promotes the proliferation and metastasis of glioma cells via the phosphatidylinositol 3 kinase-matrix metalloproteinase 2 (AKT-MMP2) pathway. Ann Transl Med 2021;9(8):659.

41. Liang Z, Li X, Chen J, Cai H, Zhang L, Li C, Tong J, Hu W. PRC1 promotes cell proliferation and cell cycle progression by regulating p21/p27-pRB family molecules and FAK-paxillin pathway in non-small cell lung cancer. Transl Cancer Res 2019;8(5):2059–2072.

42. Xiong Y, Lu J, Fang Q, Lu Y, Xie C, Wu H, Yin Z. UBE2C functions as a potential oncogene by enhancing cell proliferation, migration, invasion, and drug resistance in hepatocellular carcinoma cells. Biosci Rep 2019;39(4):BSR20182384.

43. Xu L, Yu W, Xiao H, Lin K. BIRC5 is a prognostic biomarker associated with tumor immune cell infiltration. Sci Rep 2021;11(1):390.

44. Mara JN, Zhou LT, Larmore M, Johnson B, Ayiku R, Amargant F, Pritchard MT, Duncan FE. Ovulation and ovarian wound healing are impaired with advanced reproductive age. Aging (Albany NY) 2020;12(10):9686–9713.

45. Florin L, Knebel J, Zigrino P, Vonderstrass B, Mauch C, Schorpp-Kistner M, Szabowski A, Angel P. Delayed wound healing and epidermal hyperproliferation in mice lacking JunB in the skin. J Invest Dermatol 2006;126(4):902–911.

46. Wu M, Melichian DS, de la Garza M, Gruner K, Bhattacharyya S, Barr L, Nair A, Shahrara S, Sporn PHS, Mustoe TA, Tourtellotte WG, Varga J. Essential roles for early growth response transcription factor Egr-1 in tissue fibrosis and wound healing. Am J Pathol 2009;175(3):1041–1055.

47. Martin P, Nobes CD. An early molecular component of the wound healing response in rat embryos--induction of c-fos protein in cells at the epidermal wound margin. Mech Dev 1992;38(3):209–215.

48. Yue C, Guo Z, Luo Y, Yuan J, Wan X, Mo Z. c-Jun Overexpression Accelerates Wound Healing in Diabetic Rats by Human Umbilical Cord-Derived Mesenchymal Stem Cells. Stem Cells Int 2020;2020:7430968.

49. Sherman BT, Hao M, Qiu J, Jiao X, Baseler MW, Lane HC, Imamichi T, Chang W. DAVID: a web server for functional enrichment analysis and functional annotation of gene lists (2021 update). Nucleic Acids Res 2022;50(W1):W216–221.

50. Huang DW, Sherman BT, Lempicki RA. Systematic and integrative analysis of large gene lists using DAVID bioinformatics resources. Nat Protoc 2009;4(1):44–57.

51. The Genotype-Tissue Expression (GTEx) project. Nat Genet 2013;45(6):580–585.

52. Toms D, Pan B, Li J. Endocrine Regulation in the Ovary by MicroRNA during the Estrous Cycle. Front Endocrinol (Lausanne) 2017;8:378.

53. Kubota K, Cui W, Dhakal P, Wolfe MW, Rumi MAK, Vivian JL, Roby KF, Soares MJ. Rethinking progesterone regulation of female reproductive cyclicity. Proc Natl Acad Sci U S A 2016;113(15):4212–4217.

54. Hwang B, Lee JH, Bang D. Single-cell RNA sequencing technologies and bioinformatics pipelines. Exp Mol Med 2018;50(8):1–14.

55. Deb S, Campbell BK, Clewes JS, Pincott-Allen C, Raine-Fenning NJ. Intracycle variation in number of antral follicles stratified by size and in endocrine markers of ovarian reserve in women with normal ovulatory menstrual cycles. Ultrasound Obstet Gynecol 2013;41(2):216–222.

56. Richards JS. Maturation of ovarian follicles: actions and interactions of pituitary and ovarian hormones on follicular cell differentiation. Physiol Rev 1980;60(1):51–89.

57. Lazzaro MA, Pépin D, Pescador N, Murphy BD, Vanderhyden BC, Picketts DJ. The imitation switch protein SNF2L regulates steroidogenic acute regulatory protein expression during terminal differentiation of ovarian granulosa cells. Mol Endocrinol 2006;20(10):2406–2417.

58. Irving-Rodgers HF, Harland ML, Sullivan TR, Rodgers RJ. Studies of granulosa cell maturation in dominant and subordinate bovine follicles: novel extracellular matrix focimatrix is co-ordinately regulated with cholesterol side-chain cleavage CYP11A1. Reproduction 2009;137(5):825–834.

59. Wang S, Liu W, Pang X, Dai S, Liu G. The Mechanism of Melatonin and Its Receptor MT2 Involved in the Development of Bovine Granulosa Cells. Int J Mol Sci 2018;19(7). doi:10.3390/ijms19072028.

60. Wahlberg P, Nylander A, Ahlskog N, Liu K, Ny T. Expression and localization of the serine proteases high-temperature requirement factor A1, serine protease 23, and serine protease 35 in the mouse ovary. Endocrinology 2008;149(10):5070–5077.

61. Lussier JG, Diouf MN, Lévesque V, Sirois J, Ndiaye K. Gene expression profiling of upregulated mRNAs in granulosa cells of bovine ovulatory follicles following stimulation with hCG. Reprod Biol Endocrinol 2017;15(1):88.

62. Sayasith K, Lussier J, Sirois J. Molecular characterization and transcriptional regulation of a disintegrin and metalloproteinase with thrombospondin motif 1 (ADAMTS1) in bovine preovulatory follicles. Endocrinology 2013;154(8):2857–2869.

63. Diaz FJ, Wigglesworth K, Eppig JJ. Oocytes determine cumulus cell lineage in mouse ovarian follicles. Journal of Cell Science 2007;120(8):1330–1340.

64. Duncan WC. The inadequate corpus luteum. Reprod Fertil 2021;2(1):C1–C7.

65. Noguchi M, Hirata M, Kawaguchi H, Tanimoto A. Corpus luteum Regression Induced by Prostaglandin F(2α) in Microminipigs During the Normal Estrous Cycle. In Vivo 2017;31(6):1097–1101.

66. Zhu R, Zou S-T, Wan J-M, Li W, Li X-L, Zhu W. BTG1 inhibits breast cancer cell growth through induction of cell cycle arrest and apoptosis. Oncol Rep 2013;30(5):2137–2144.

67. Ryan KJ, Petro Z. Steroid Biosynthesis by Human Ovarian Granulosa and Thecal Cells. The Journal of Clinical Endocrinology & Metabolism 1966;26(1):46–52.

68. Hummitzsch K, Hatzirodos N, Macpherson AM, Schwartz J, Rodgers RJ, Irving-Rodgers HF. Transcriptome analyses of ovarian stroma: tunica albuginea, interstitium and theca interna. Reproduction 2019;157(6):545–565.

69. Grosdemouge I, Bachelot A, Lucas A, Baran N, Kelly PA, Binart N. Effects of deletion of the prolactin receptor on ovarian gene expression. Reprod Biol Endocrinol 2003;1:12.

70. Sheng X, Zhou J, Kang N, Liu W, Yu L, Zhang Z, Zhang Y, Yue Q, Yang Q, Zhang X, Li C, Yan G, Sun H. Temporal and spatial dynamics mapping reveals follicle development regulated by different stromal cell populations. bioRxiv 2022. doi:10.1101/2022.03.04.480328.

71. Kinnear HM, Tomaszewski CE, Chang FL, Moravek MB, Xu M, Padmanabhan V, Shikanov A. The ovarian stroma as a new frontier. Reproduction 2020;160(3):R25–R39.

72. Duffy DM, Ko C, Jo M, Brannstrom M, Curry TE. Ovulation: Parallels With Inflammatory Processes. Endocr Rev 2019;40(2):369–416.

73. Hartanti MD, Hummitzsch K, Irving-Rodgers HF, Bonner WM, Copping KJ, Anderson RA, McMillen IC, Perry VEA, Rodgers RJ. Morphometric and gene expression analyses of stromal expansion during development of the bovine fetal ovary. Reprod Fertil Dev 2019;31(3):482–495.

74. Xu F, Stouffer RL, Müller J, Hennebold JD, Wright JW, Bahar A, Leder G, Peters M, Thorne M, Sims M, Wintermantel T, Lindenthal B. Dynamics of the transcriptome in the primate ovulatory follicle. Mol Hum Reprod 2011;17(3):152–165.

75. Kenngott RA-M, Sauer U, Vermehren M, Sinowatz F. Expression of Intermediate Filaments and Germ Cell Markers in the Developing Bovine Ovary: An Immunohistochemical and Laser-Assisted Microdissection Study. Cells Tissues Organs 2014;200(2):153–170.

76. Lin Y-T, Chen J-S, Wu M-H, Hsieh I-S, Liang C-H, Hsu C-L, Hong T-M, Chen Y-L. Galectin-1 accelerates wound healing by regulating the neuropilin-1/Smad3/NOX4 pathway and ROS production in myofibroblasts. J Invest Dermatol 2015;135(1):258–268.

77. Okada Y, Saika S, Hashizume N, Kobata S, Yamanaka O, Ohnishi Y, Senba E. Expression of fos family and jun family proto-oncogenes during corneal epithelial wound healing. Curr Eye Res 1996;15(8):824–832.

78. Wijayarathna R, de Kretser DM. Activins in reproductive biology and beyond. Hum Reprod Update 2016;22(3):342–357.

79. Li S-H, Lin M-H, Hwu Y-M, Lu C-H, Yeh L-Y, Chen Y-J, Lee RK-K. Correlation of cumulus gene expression of GJA1, PRSS35, PTX3, and SERPINE2 with oocyte maturation, fertilization, and embryo development. Reprod Biol Endocrinol 2015;13:93.

80. Zhang M, Su Y-Q, Sugiura K, Xia G, Eppig JJ. Granulosa cell ligand NPPC and its receptor NPR2 maintain meiotic arrest in mouse oocytes. Science 2010;330(6002):366–369.

81. Xi G, An L, Wang W, Hao J, Yang Q, Ma L, Lu J, Wang Y, Wang W, Zhao W, Liu J, Yang M, Wang X, Zhang Z, Zhang C, Chu M, Yue Y, Yao F, Zhang M, Tian J. The mRNA-destabilizing protein Tristetraprolin targets “meiosis arrester” Nppc mRNA in mammalian preovulatory follicles. Proc Natl Acad Sci U S A 2021;118(22). doi:10.1073/pnas.2018345118.

82. Akaiwa M, Fukui E, Matsumoto H. Tubulointerstitial nephritis antigen-like 1 deficiency alleviates age-dependent depressed ovulation associated with ovarian collagen deposition in mice. Reprod Med Biol 2020;19(1):50–57.

83. Kim Y-S, Hwan JD, Bae S, Bae D-H, Shick WA. Identification of differentially expressed genes using an annealing control primer system in stage III serous ovarian carcinoma. BMC Cancer 2010;10:576.

84. Chang H-M, Leung PCK. Physiological roles of activins in the human ovary. Journal of Bio-X Research 2018;1(3). Available at: https://journals.lww.com/jbioxresearch/Fulltext/2018/12000/Physiological_roles_of_activins_in_the_human_ovary.2.aspx.

85. O’Connell AR, McNatty KP, Hurst PR, Spencer TE, Bazer FW, Reader KL, Johnstone PD, Davis GH, Juengel JL. Activin A and follistatin during the oestrous cycle and early pregnancy in ewes. J Endocrinol 2016;228(3):193–203.

86. Celik O, Celik N, Ugur K, Hatirnaz S, Celik S, Muderris II, Yavuzkir S, Sahin i, Yardim M, Aydin S. Nppc/Npr2/cGMP signaling cascade maintains oocyte developmental capacity. Cell Mol Biol (Noisy-le-grand) 2019;65(4):83–89.

87. Celik O, Celik N, Gungor S, Haberal ET, Aydin S. Selective Regulation of Oocyte Meiotic Events Enhances Progress in Fertility Preservation Methods. Biochem Insights 2015;8:11–21.

88. Meinsohn M-C, Morin F, Bertolin K, Duggavathi R, Schoonjans K, Murphy BD. The Orphan Nuclear Receptor Liver Homolog Receptor-1 (Nr5a2) Regulates Ovarian Granulosa Cell Proliferation. J Endocr Soc 2018;2(1):24–41.

89. Hodgkinson K, Forrest LA, Vuong N, Garson K, Djordjevic B, Vanderhyden BC. GREB1 is an estrogen receptor-regulated tumour promoter that is frequently expressed in ovarian cancer. Oncogene 2018;37(44):5873–5886.

90. Terman BI, Dougher-Vermazen M, Carrion ME, Dimitrov D, Armellino DC, Gospodarowicz D, Böhlen P. Identification of the KDR tyrosine kinase as a receptor for vascular endothelial cell growth factor. Biochem Biophys Res Commun 1992;187(3):1579–1586.

91. Parikh A, Lee C, Joseph P, Marchini S, Baccarini A, Kolev V, Romualdi C, Fruscio R, Shah H, Wang F, Mullokandov G, Fishman D, D’Incalci M, Rahaman J, Kalir T, Redline RW, Brown BD, Narla G, DiFeo A. microRNA-181a has a critical role in ovarian cancer progression through the regulation of the epithelial-mesenchymal transition. Nat Commun 2014;5:2977.

92. Galvagni F, Nardi F, Spiga O, Trezza A, Tarticchio G, Pellicani R, Andreuzzi E, Caldi E, Toti P, Tosi GM, Santucci A, Iozzo RV, Mongiat M, Orlandini M. Dissecting the CD93-Multimerin 2 interaction involved in cell adhesion and migration of the activated endothelium. Matrix Biol 2017;64:112–127.

93. Richards JS, Fan H-Y, Liu Z, Tsoi M, Laguë M-N, Boyer A, Boerboom D. Either Kras activation or Pten loss similarly enhance the dominant-stable CTNNB1-induced genetic program to promote granulosa cell tumor development in the ovary and testis. Oncogene 2012;31(12):1504–1520.

94. Zhou Q, Wan M, Wei Q, Song Q, Xiong L, Huo J, Huang J. Expression, Regulation, and Functional Characterization of FST Gene in Porcine Granulosa Cells. Anim Biotechnol 2016;27(4):295–302.

95. Bambino K, Lacko LA, Hajjar KA, Stuhlmann H. Epidermal growth factor-like domain 7 is a marker of the endothelial lineage and active angiogenesis. Genesis 2014;52(7):657–670.

96. He W, Gauri M, Li T, Wang R, Lin S-X. Current knowledge of the multifunctional 17β-hydroxysteroid dehydrogenase type 1 (HSD17B1). Gene 2016;588(1):54–61.

97. Oksjoki S, Sallinen S, Vuorio E, Anttila L. Cyclic expression of mRNA transcripts for connective tissue components in the mouse ovary. Mol Hum Reprod 1999;5(9):803–808.

98. Park S, DiMaio TA, Scheef EA, Sorenson CM, Sheibani N. PECAM-1 regulates proangiogenic properties of endothelial cells through modulation of cell-cell and cell-matrix interactions. Am J Physiol Cell Physiol 2010;299(6):C1468–1484.

99. Wigglesworth K, Lee K-B, Emori C, Sugiura K, Eppig JJ. Transcriptomic diversification of developing cumulus and mural granulosa cells in mouse ovarian follicles. Biol Reprod 2015;92(1):23.

100. Varankar SS, More M, Abraham A, Pansare K, Kumar B, Narayanan NJ, Jolly MK, Mali AM, Bapat SA. Functional balance between Tcf21-Slug defines cellular plasticity and migratory modalities in high grade serous ovarian cancer cell lines. Carcinogenesis 2020;41(4):515–526.

101. Herr D, Fraser HM, Konrad R, Holzheu I, Kreienberg R, Wulff C. Human chorionic gonadotropin controls luteal vascular permeability via vascular endothelial growth factor by down-regulation of a cascade of adhesion proteins. Fertil Steril 2013;99(6):1749–1758.

102. Angelos MG, Abrahante JE, Blum RH, Kaufman DS. Single Cell Resolution of Human Hematoendothelial Cells Defines Transcriptional Signatures of Hemogenic Endothelium. Stem Cells 2018;36(2):206–217.

103. Bédard J, Brûlé S, Price CA, Silversides DW, Lussier JG. Serine protease inhibitor-E2 (SERPINE2) is differentially expressed in granulosa cells of dominant follicle in cattle. Mol Reprod Dev 2003;64(2):152–165.

104. Sleer LS, Taylor CC. Cell-type localization of platelet-derived growth factors and receptors in the postnatal rat ovary and follicle. Biol Reprod 2007;76(3):379–390.

105. de Vries C, Escobedo JA, Ueno H, Houck K, Ferrara N, Williams LT. The fms-like tyrosine kinase, a receptor for vascular endothelial growth factor. Science 1992;255(5047):989–991.

106. Wu Y, Lin J, Li X, Han B, Wang L, Liu M, Huang J. Transcriptome profile of one-month-old lambs’ granulosa cells after superstimulation. Asian-Australas J Anim Sci 2017;30(1):20–33.

107. Galvagni F, Nardi F, Maida M, Bernardini G, Vannuccini S, Petraglia F, Santucci A, Orlandini M. CD93 and dystroglycan cooperation in human endothelial cell adhesion and migration adhesion and migration. Oncotarget 2016;7(9):10090–10103.

108. Sterzyńska K, Klejewski A, Wojtowicz K, Świerczewska M, Andrzejewska M, Rusek D, Sobkowski M, Kędzia W, Brązert J, Nowicki M, Januchowski R. The Role of Matrix Gla Protein (MGP) Expression in Paclitaxel and Topotecan Resistant Ovarian Cancer Cell Lines. Int J Mol Sci 2018;19(10). doi:10.3390/ijms19102901.

109. Kalucka J, de Rooij LPMH, Goveia J, Rohlenova K, Dumas SJ, Meta E, Conchinha NV, Taverna F, Teuwen L-A, Veys K, García-Caballero M, Khan S, Geldhof V, Sokol L, Chen R, Treps L, Borri M, de Zeeuw P, Dubois C, Karakach TK, Falkenberg KD, Parys M, Yin X, Vinckier S, Du Y, Fenton RA, Schoonjans L, Dewerchin M, Eelen G, Thienpont B, Lin L, Bolund L, Li X, Luo Y, Carmeliet P. Single-Cell Transcriptome Atlas of Murine Endothelial Cells. Cell 2020;180(4):764–779.e20.

110. Cochain C, Vafadarnejad E, Arampatzi P, Pelisek J, Winkels H, Ley K, Wolf D, Saliba A-E, Zernecke A. Single-Cell RNA-Seq Reveals the Transcriptional Landscape and Heterogeneity of Aortic Macrophages in Murine Atherosclerosis. Circ Res 2018;122(12):1661–1674.

111. Carpenter AR, Becknell MB, Ching CB, Cuaresma EJ, Chen X, Hains DS, McHugh KM. Uroplakin 1b is critical in urinary tract development and urothelial differentiation and homeostasis. Kidney Int 2016;89(3):612–624.

112. Glowacka WK, Alberts P, Ouchida R, Wang J-Y, Rotin D. LAPTM5 protein is a positive regulator of proinflammatory signaling pathways in macrophages. J Biol Chem 2012;287(33):27691–27702.

113. Paulini F, Melo EO. The role of oocyte-secreted factors GDF9 and BMP15 in follicular development and oogenesis. Reprod Domest Anim 2011;46(2):354–361.

114. Rudat C, Grieskamp T, Röhr C, Airik R, Wrede C, Hegermann J, Herrmann BG, Schuster-Gossler K, Kispert A. Upk3b is dispensable for development and integrity of urothelium and mesothelium. PLoS One 2014;9(11):e112112.

115. Yang P, Wu Q, Sun L, Fang P, Liu L, Ji Y, Park J-Y, Qin X, Yang X, Wang H. Adaptive Immune Response Signaling Is Suppressed in Ly6C(high) Monocyte but Upregulated in Monocyte Subsets of ApoE (-/-) Mice - Functional Implication in Atherosclerosis. Front Immunol 2021;12:809208.

116. Tanaka M, Kihara M, Hennebold JD, Eppig JJ, Viveiros MM, Emery BR, Carrell DT, Kirkman NJ, Meczekalski B, Zhou J, Bondy CA, Becker M, Schultz RM, Misteli T, De La Fuente R, King GJ, Adashi EY. H1FOO is coupled to the initiation of oocytic growth. Biol Reprod 2005;72(1):135–142.

117. Sewgobind NV, Albers S, Pieters RJ. Functions and Inhibition of Galectin-7, an Emerging Target in Cellular Pathophysiology. Biomolecules 2021;11(11). doi:10.3390/biom11111720.

118. Liu X, Morency E, Li T, Qin H, Zhang X, Zhang X, Coonrod S. Role for PADI6 in securing the mRNA-MSY2 complex to the oocyte cytoplasmic lattices. Cell Cycle 2017;16(4):360–366.

119. Auersperg N. The stem-cell profile of ovarian surface epithelium is reproduced in the oviductal fimbriae, with increased stem-cell marker density in distal parts of the fimbriae. Int J Gynecol Pathol 2013;32(5):444–453.

120. Sweet RA, Nickerson KM, Cullen JL, Wang Y, Shlomchik MJ. B Cell-Extrinsic Myd88 and Fcer1g Negatively Regulate Autoreactive and Normal B Cell Immune Responses. J Immunol 2017;199(3):885–893.

121. Bebbere D, Masala L, Albertini DF, Ledda S. The subcortical maternal complex: multiple functions for one biological structure? J Assist Reprod Genet 2016;33(11):1431–1438.

122. Iwata M, Takebayashi T, Ohta H, Alcalde RE, Itano Y, Matsumura T. Zinc accumulation and metallothionein gene expression in the proliferating epidermis during wound healing in mouse skin. Histochem Cell Biol 1999;112(4):283–290.

123. Gava N, L Clarke C, Bye C, Byth K, deFazio A. Global gene expression profiles of ovarian surface epithelial cells in vivo. J Mol Endocrinol 2008;40(6):281–296.

124. Paillisson A, Dadé S, Callebaut I, Bontoux M, Dalbiès-Tran R, Vaiman D, Monget P. Identification, characterization and metagenome analysis of oocyte-specific genes organized in clusters in the mouse genome. BMC Genomics 2005;6:76.

125. Higashi T, Tokuda S, Kitajiri S, Masuda S, Nakamura H, Oda Y, Furuse M. Analysis of the “angulin” proteins LSR, ILDR1 and ILDR2--tricellulin recruitment, epithelial barrier function and implication in deafness pathogenesis. J Cell Sci 2013;126(Pt 4):966–977.

126. Stables MJ, Shah S, Camon EB, Lovering RC, Newson J, Bystrom J, Farrow S, Gilroy DW. Transcriptomic analyses of murine resolution-phase macrophages. Blood 2011;118(26):e192–208.

127. Rajkovic A, Lee JH, Yan C, Matzuk MM. The ret finger protein-like 4 gene, Rfpl4, encodes a putative E3 ubiquitin-protein ligase expressed in adult germ cells. Mech Dev 2002;112(1–2):173–177.

128. Sontheimer RD, Racila E, Racila DM. C1q: its functions within the innate and adaptive immune responses and its role in lupus autoimmunity. J Invest Dermatol 2005;125(1):14–23.

129. Cai C, Tamai K, Molyneaux K. KHDC1B is a novel CPEB binding partner specifically expressed in mouse oocytes and early embryos. Mol Biol Cell 2010;21(18):3137–3148.

130. Suto F, Murakami Y, Nakamura F, Goshima Y, Fujisawa H. Identification and characterization of a novel mouse plexin, plexin-A4. Mech Dev 2003;120(3):385–396.

131. Choi Y, Yuan D, Rajkovic A. Germ cell-specific transcriptional regulator sohlh2 is essential for early mouse folliculogenesis and oocyte-specific gene expression. Biol Reprod 2008;79(6):1176–1182.

132. Sakai N, Bain G, Furuichi K, Iwata Y, Nakamura M, Hara A, Kitajima S, Sagara A, Miyake T, Toyama T, Sato K, Nakagawa S, Shimizu M, Kaneko S, Wada T. The involvement of autotaxin in renal interstitial fibrosis through regulation of fibroblast functions and induction of vascular leakage. Sci Rep 2019;9(1):7414.

133. Puttabyatappa M, Guo X, Dou J, Dumesic D, Bakulski KM, Padmanabhan V. Developmental Programming: Sheep Granulosa and Theca Cell-Specific Transcriptional Regulation by Prenatal Testosterone. Endocrinology 2020;161(8). doi:10.1210/endocr/bqaa094.

134. Singh H, Li Y, Fuller PJ, Harrison C, Rao J, Stephens AN, Nie G. HtrA3 Is Downregulated in Cancer Cell Lines and Significantly Reduced in Primary Serous and Granulosa Cell Ovarian Tumors. J Cancer 2013;4(2):152–164.

135. Luo F, Jia R, Ying S, Wang Z, Wang F. Analysis of genes that influence sheep follicular development by different nutrition levels during the luteal phase using expression profiling. Anim Genet 2016;47(3):354–364.

136. Cheng Y, Kawamura K, Deguchi M, Takae S, Mulders SM, Hsueh AJW. Intraovarian thrombin and activated protein C signaling system regulates steroidogenesis during the periovulatory period. Mol Endocrinol 2012;26(2):331–340.

137. Hatzirodos N, Glister C, Hummitzsch K, Irving-Rodgers HF, Knight PG, Rodgers RJ. Transcriptomal profiling of bovine ovarian granulosa and theca interna cells in primary culture in comparison with their in vivo counterparts. PLoS One 2017;12(3):e0173391.

138. Lawrence JB, Oxvig C, Overgaard MT, Sottrup-Jensen L, Gleich GJ, Hays LG, Yates JR 3rd, Conover CA. The insulin-like growth factor (IGF)-dependent IGF binding protein-4 protease secreted by human fibroblasts is pregnancy-associated plasma protein-A. Proc Natl Acad Sci U S A 1999;96(6):3149–3153.

139. Hu S, Tamada K, Ni J, Vincenz C, Chen L. Characterization of TNFRSF19, a novel member of the tumor necrosis factor receptor superfamily. Genomics 1999;62(1):103–107.

140. Reverchon M, Bertoldo MJ, Ramé C, Froment P, Dupont J. CHEMERIN (RARRES2) decreases in vitro granulosa cell steroidogenesis and blocks oocyte meiotic progression in bovine species. Biol Reprod 2014;90(5):102.

141. Guerrero-Juarez CF, Dedhia PH, Jin S, Ruiz-Vega R, Ma D, Liu Y, Yamaga K, Shestova O, Gay DL, Yang Z, Kessenbrock K, Nie Q, Pear WS, Cotsarelis G, Plikus MV. Single-cell analysis reveals fibroblast heterogeneity and myeloid-derived adipocyte progenitors in murine skin wounds. Nat Commun 2019;10(1):650.

142. Veldhuis JD. Interactions among endocrine control systems in the regulation of ovarian function. Clin Biochem 1981;14(5):252–257.

143. Xie T, Wang Y, Deng N, Huang G, Taghavifar F, Geng Y, Liu N, Kulur V, Yao C, Chen P, Liu Z, Stripp B, Tang J, Liang J, Noble PW, Jiang D. Single-Cell Deconvolution of Fibroblast Heterogeneity in Mouse Pulmonary Fibrosis. Cell Rep 2018;22(13):3625–3640.

144. Kim J-M, Song K-S, Xu B, Wang T. Role of potassium channels in female reproductive system. Obstet Gynecol Sci 2020;63(5):565–576.

145. Katoh D, Kozuka Y, Noro A, Ogawa T, Imanaka-Yoshida K, Yoshida T. Tenascin-C Induces Phenotypic Changes in Fibroblasts to Myofibroblasts with High Contractility through the Integrin αvβ1/Transforming Growth Factor β/SMAD Signaling Axis in Human Breast Cancer. Am J Pathol 2020;190(10):2123–2135.

146. Machelon V, Gaudin F, Camilleri-Broët S, Nasreddine S, Bouchet-Delbos L, Pujade-Lauraine E, Alexandre J, Gladieff L, Arenzana-Seisdedos F, Emilie D, Prévot S, Broët P, Balabanian K. CXCL12 expression by healthy and malignant ovarian epithelial cells. BMC Cancer 2011;11:97.

147. Popple A, Durrant LG, Spendlove I, Rolland P, Scott IV, Deen S, Ramage JM. The chemokine, CXCL12, is an independent predictor of poor survival in ovarian cancer. Br J Cancer 2012;106(7):1306–1313.

148. Mitchell TS, Bradley J, Robinson GS, Shima DT, Ng Y-S. RGS5 expression is a quantitative measure of pericyte coverage of blood vessels. Angiogenesis 2008;11(2):141–151.

149. Nagaraja AK, Middlebrook BS, Rajanahally S, Myers M, Li Q, Matzuk MM, Pangas SA. Defective gonadotropin-dependent ovarian folliculogenesis and granulosa cell gene expression in inhibin-deficient mice. Endocrinology 2010;151(10):4994–5006.

150. Kofler NM, Cuervo H, Uh MK, Murtomäki A, Kitajewski J. Combined deficiency of Notch1 and Notch3 causes pericyte dysfunction, models CADASIL, and results in arteriovenous malformations. Sci Rep 2015;5:16449.

151. Maguchi M, Nishida W, Kohara K, Kuwano A, Kondo I, Hiwada K. Molecular cloning and gene mapping of human basic and acidic calponins. Biochem Biophys Res Commun 1995;217(1):238–244.

152. Camoretti-Mercado B, Forsythe SM, LeBeau MM, Espinosa R 3rd, Vieira JE, Halayko AJ, Willadsen S, Kurtz B, Ober C, Evans GA, Thweatt R, Shapiro S, Niu Q, Qin Y, Padrid PA, Solway J. Expression and cytogenetic localization of the human SM22 gene (TAGLN). Genomics 1998;49(3):452–457.

153. Palandri A, L’hôte D, Cohen-Tannoudji J, Tricoire H, Monnier V. Frataxin inactivation leads to steroid deficiency in flies and human ovarian cells. Hum Mol Genet 2015;24(9):2615–2626.

154. Krawczyk KK, Skovsted GF, Perisic L, Dreier R, Berg JO, Hedin U, Rippe C, Swärd K. Expression of endothelin type B receptors (EDNRB) on smooth muscle cells is controlled by MKL2, ternary complex factors, and actin dynamics. Am J Physiol Cell Physiol 2018;315(6):C873–C884.

155. Conley CA, Fritz-Six KL, Almenar-Queralt A, Fowler VM. Leiomodins: larger members of the tropomodulin (Tmod) gene family. Genomics 2001;73(2):127–139.

156. O’Shannessy DJ, Somers EB, Wang L-C, Wang H, Hsu R. Expression of folate receptors alpha and beta in normal and cancerous gynecologic tissues: correlation of expression of the beta isoform with macrophage markers. J Ovarian Res 2015;8:29.

157. Weng J, Liao M, Zou S, Bao J, Zhou J, Qu L, Feng R, Feng X, Zhao Z, Jing Z. Downregulation of FHL1 expression in thoracic aortic dissection: implications in aortic wall remodeling and pathogenesis of thoracic aortic dissection. Ann Vasc Surg 2011;25(2):240–247.

158. Hayashi K-G, Ushizawa K, Hosoe M, Takahashi T. Differential gene expression of serine protease inhibitors in bovine ovarian follicle: possible involvement in follicular growth and atresia. Reprod Biol Endocrinol 2011;9:72.

159. Li L, Mo H, Zhang J, Zhou Y, Peng X, Luo X. The Role of Heat Shock Protein 90B1 in Patients with Polycystic Ovary Syndrome. PLoS One 2016;11(4):e0152837.

160. Chen AQ, Wang ZG, Xu ZR, Yu SD, Yang ZG. Analysis of gene expression in granulosa cells of ovine antral growing follicles using suppressive subtractive hybridization. Anim Reprod Sci 2009;115(1–4):39–48.

161. Lee JH, Berger JM. Cell Cycle-Dependent Control and Roles of DNA Topoisomerase II. Genes (Basel) 2019;10(11). doi:10.3390/genes10110859.

162. Tsuji T, Kiyosu C, Akiyama K, Kunieda T. CNP/NPR2 signaling maintains oocyte meiotic arrest in early antral follicles and is suppressed by EGFR-mediated signaling in preovulatory follicles. Mol Reprod Dev 2012;79(11):795–802.

163. Xi G, Wang W, Fazlani SA, Yao F, Yang M, Hao J, An L, Tian J. C-type natriuretic peptide enhances mouse preantral follicle growth. Reproduction 2019;157(5):445–455.

164. Piersanti RL, Santos JEP, Sheldon IM, Bromfield JJ. Lipopolysaccharide and tumor necrosis factor-alpha alter gene expression of oocytes and cumulus cells during bovine in vitro maturation. Mol Reprod Dev 2019;86(12):1909–1920.

165. Alfieri C, Chang L, Zhang Z, Yang J, Maslen S, Skehel M, Barford D. Molecular basis of APC/C regulation by the spindle assembly checkpoint. Nature 2016;536(7617):431–436.

166. Rovani MT, Gasperin BG, Ilha GF, Ferreira R, Bohrer RC, Duggavathi R, Bordignon V, Gonçalves PBD. Expression and molecular consequences of inhibition of estrogen receptors in granulosa cells of bovine follicles. J Ovarian Res 2014;7:96.

167. Hatzirodos N, Irving-Rodgers HF, Hummitzsch K, Rodgers RJ. Transcriptome profiling of the theca interna from bovine ovarian follicles during atresia. PLoS One 2014;9(6):e99706.

168. Landry DA, Rossi-Perazza L, Lafontaine S, Sirard M-A. Expression of atresia biomarkers in granulosa cells after ovarian stimulation in heifers. Reproduction 2018;156(3):239–248.

169. Diaz FJ, Sugiura K, Eppig JJ. Regulation of Pcsk6 expression during the preantral to antral follicle transition in mice: opposing roles of FSH and oocytes. Biol Reprod 2008;78(1):176–183.

170. Blanchard JM. Cyclin A2 transcriptional regulation: modulation of cell cycle control at the G1/S transition by peripheral cues. Biochem Pharmacol 2000;60(8):1179–1184.

171. Gong D, Ferrell JEJ. The roles of cyclin A2, B1, and B2 in early and late mitotic events. Mol Biol Cell 2010;21(18):3149–3161.

172. Luo Y, Qiao X, Ma Y, Deng H, Xu CC, Xu L. Irisin deletion induces a decrease in growth and fertility in mice. Reprod Biol Endocrinol 2021;19(1):22.

173. Pallier C, Scaffidi P, Chopineau-Proust S, Agresti A, Nordmann P, Bianchi ME, Marechal V. Association of chromatin proteins high mobility group box (HMGB) 1 and HMGB2 with mitotic chromosomes. Mol Biol Cell 2003;14(8):3414–3426.

174. Chun SY, McGee EA, Hsu SY, Minami S, LaPolt PS, Yao HH, Bahr JM, Gougeon A, Schomberg DW, Hsueh AJ. Restricted expression of WT1 messenger ribonucleic acid in immature ovarian follicles: uniformity in mammalian and avian species and maintenance during reproductive senescence. Biol Reprod 1999;60(2):365–373.

175. Gao F, Zhang J, Wang X, Yang J, Chen D, Huff V, Liu Y-X. Wt1 functions in ovarian follicle development by regulating granulosa cell differentiation. Hum Mol Genet 2014;23(2):333–341.

176. Park M, Choi Y, Choi H, Roh J. Wilms’ tumor suppressor gene (WT1) suppresses apoptosis by transcriptionally downregulating BAX expression in immature rat granulosa cells. J Ovarian Res 2014;7:118.

177. Gassmann R, Carvalho A, Henzing AJ, Ruchaud S, Hudson DF, Honda R, Nigg EA, Gerloff DL, Earnshaw WC. Borealin: a novel chromosomal passenger required for stability of the bipolar mitotic spindle. J Cell Biol 2004;166(2):179–191.

178. Bertolin K, Meinsohn M-C, Suzuki J, Gossen J, Schoonjans K, Duggavathi R, Murphy BD. Ovary-specific depletion of the nuclear receptor Nr5a2 compromises expansion of the cumulus oophorus but not fertilization by intracytoplasmic sperm injection. Biol Reprod 2017;96(6):1231–1243.

179. Jones MC, Zha J, Humphries MJ. Connections between the cell cycle, cell adhesion and the cytoskeleton. Philos Trans R Soc Lond B Biol Sci 2019;374(1779):20180227.

180. Bellanger S, de Gramont A, Sobczak-Thépot J. Cyclin B2 suppresses mitotic failure and DNA re-replication in human somatic cells knocked down for both cyclins B1 and B2. Oncogene 2007;26(51):7175–7184.

181. Huang Y, Sramkoski RM, Jacobberger JW. The kinetics of G2 and M transitions regulated by B cyclins. PLoS One 2013;8(12):e80861.

182. Berisha B, Rodler D, Schams D, Sinowatz F, Pfaffl MW. Prostaglandins in Superovulation Induced Bovine Follicles During the Preovulatory Period and Early Corpus Luteum. Front Endocrinol (Lausanne) 2019;10:467.

183. Kfir S, Basavaraja R, Wigoda N, Ben-Dor S, Orr I, Meidan R. Genomic profiling of bovine corpus luteum maturation. PLoS One 2018;13(3):e0194456.

184. Tamura K, Matsushita M, Endo A, Kutsukake M, Kogo H. Effect of insulin-like growth factor-binding protein 7 on steroidogenesis in granulosa cells derived from equine chorionic gonadotropin-primed immature rat ovaries. Biol Reprod 2007;77(3):485–491.

185. Duan WR, Parmer TG, Albarracin CT, Zhong L, Gibori G. PRAP, a prolactin receptor associated protein: its gene expression and regulation in the corpus luteum. Endocrinology 1997;138(8):3216–3221.

186. Hanaue M, Miwa N, Takamatsu K. Immunohistochemical Characterization of S100A6 in the Murine Ovary. Acta Histochem Cytochem 2012;45(1):9–14.

187. Hernandez-Gonzalez I, Gonzalez-Robayna I, Shimada M, Wayne CM, Ochsner SA, White L, Richards JS. Gene expression profiles of cumulus cell oocyte complexes during ovulation reveal cumulus cells express neuronal and immune-related genes: does this expand their role in the ovulation process? Mol Endocrinol 2006;20(6):1300–1321.

188. Zamberlam G, Lapointe E, Abedini A, Rico C, Godin P, Paquet M, DeMayo FJ, Boerboom D. SFRP4 Is a Negative Regulator of Ovarian Follicle Development and Female Fertility. Endocrinology 2019;160(7):1561–1572.

189. Wang SJ, Liu WJ, Wu CJ, Ma FH, Ahmad S, Liu BR, Han L, Jiang XP, Zhang SJ, Yang LG. Melatonin suppresses apoptosis and stimulates progesterone production by bovine granulosa cells via its receptors (MT1 and MT2). Theriogenology 2012;78(7):1517–1526.

190. Lee-Thacker S, Choi Y, Taniuchi I, Takarada T, Yoneda Y, Ko C, Jo M. Core Binding Factor β Expression in Ovarian Granulosa Cells Is Essential for Female Fertility. Endocrinology 2018;159(5):2094–2109.

191. Park JY, Jang H, Curry TE, Sakamoto A, Jo M. Prostate androgen-regulated mucin-like protein 1: a novel regulator of progesterone metabolism. Mol Endocrinol 2013;27(11):1871–1886.

192. Baufeld A, Koczan D, Vanselow J. Induction of altered gene expression profiles in cultured bovine granulosa cells at high cell density. Reprod Biol Endocrinol 2017;15(1):3.

193. Kapfhamer J, Waite C, Ascoli M. The Gα(q/11)-provoked induction of Akr1c18 in murine luteal cells is mediated by phospholipase C. Mol Cell Endocrinol 2018;470:179–187.

