## Supplementary figures and images for "A single cell atlas of the cycling murine ovary"

### Figure1 S1

Figure 1- supplement 1

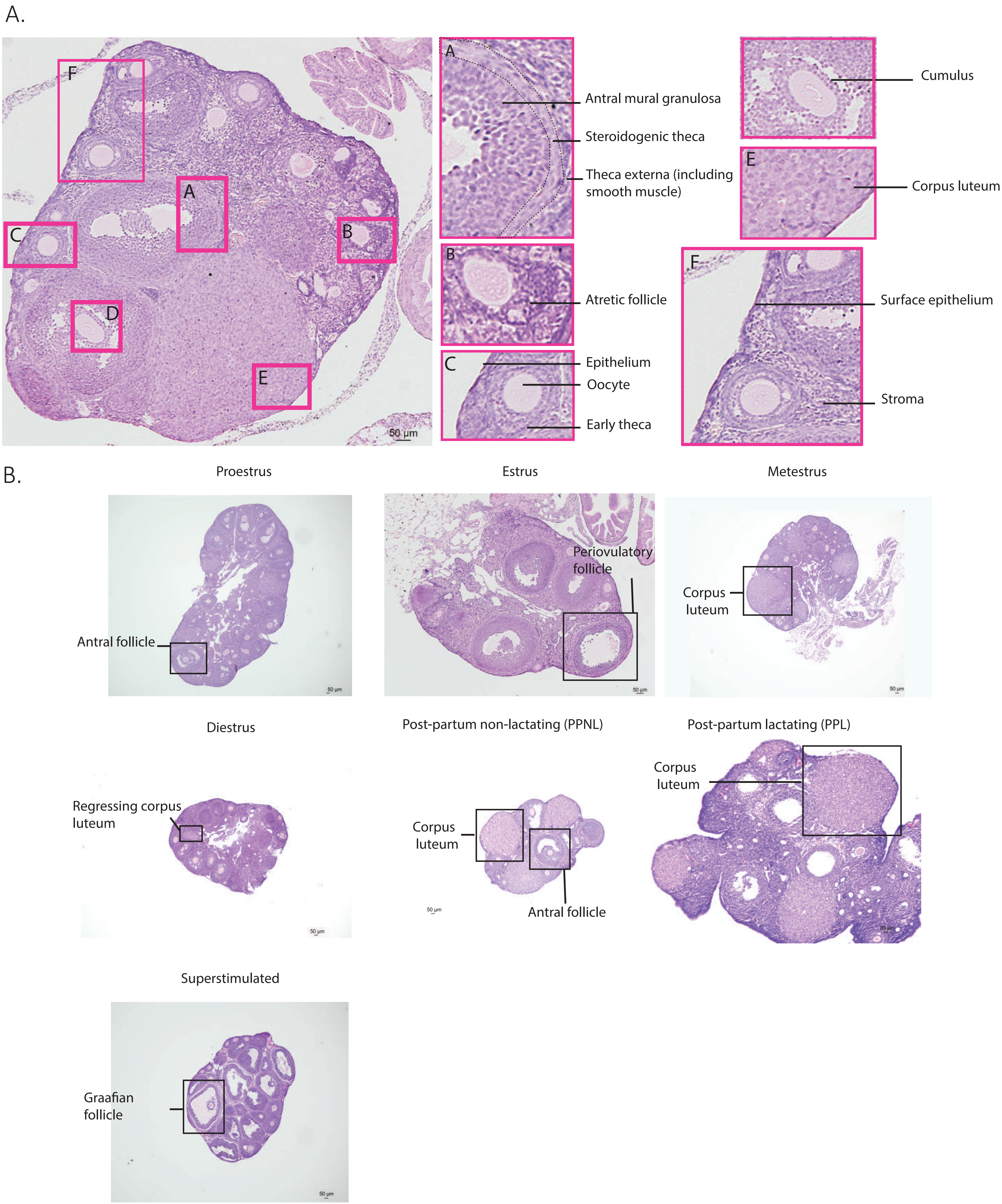

### Figure2 S1

#### Figure 2- supplement 1

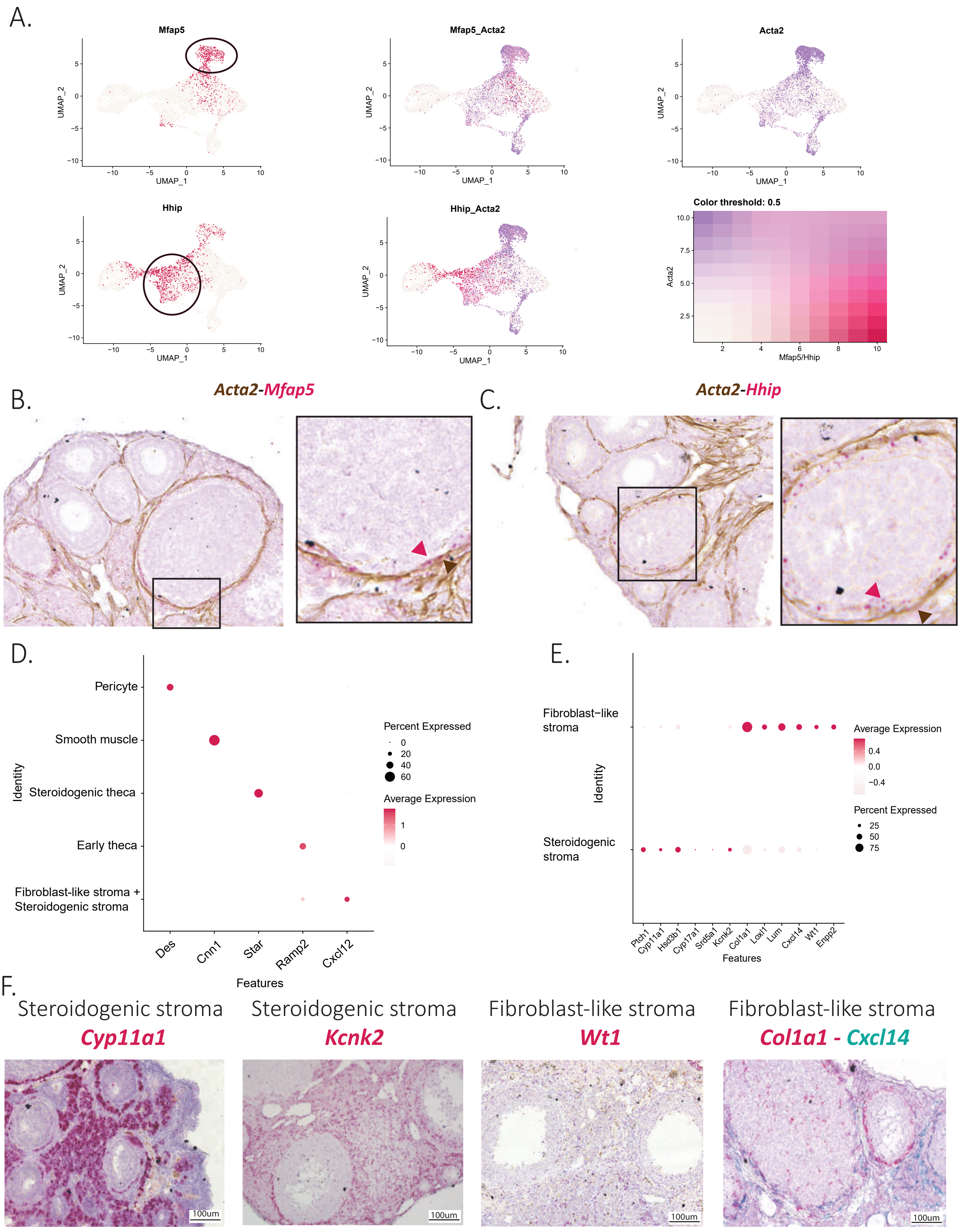

### Figure3 S1

Figure 3- supplement 1

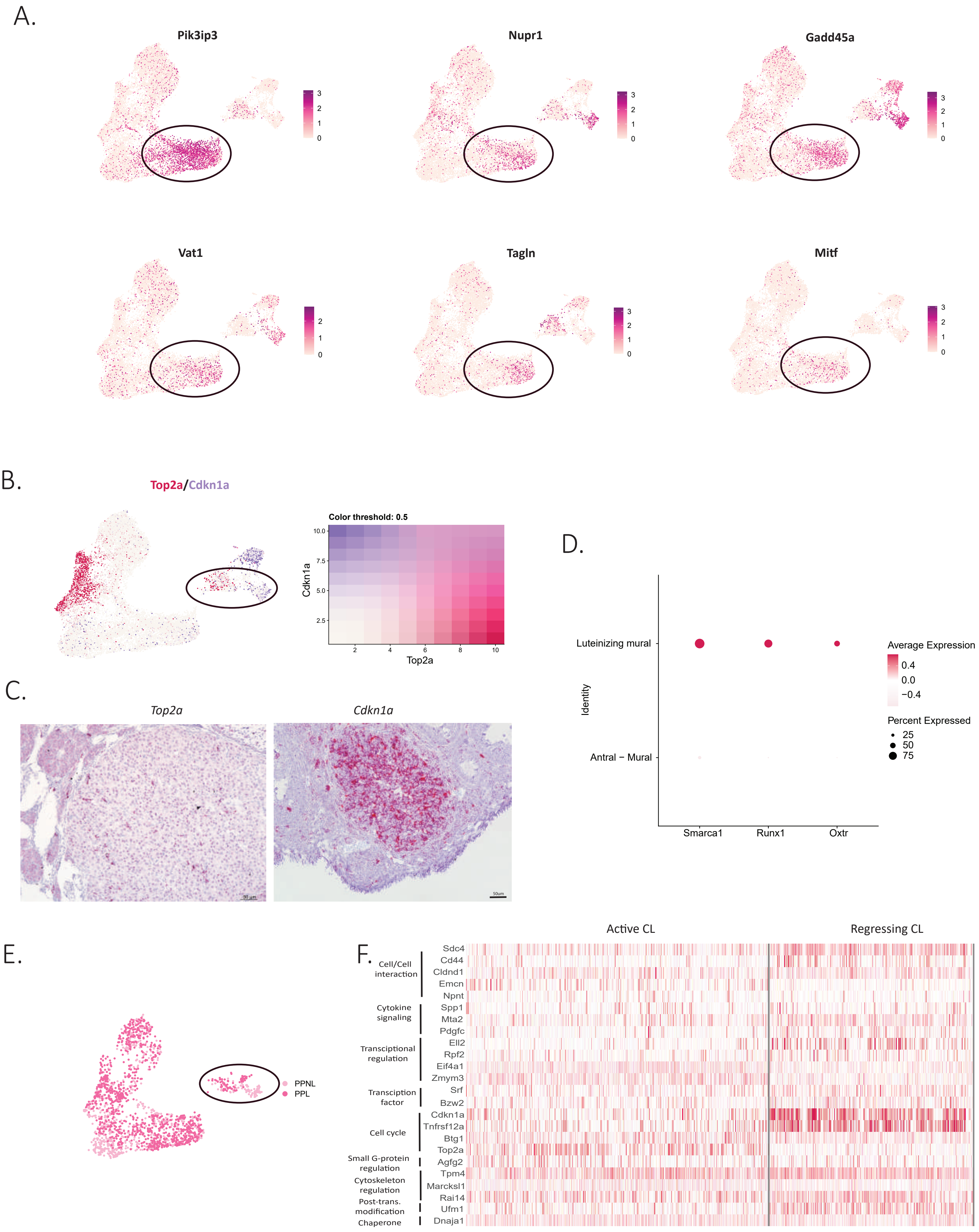

### Figure 5 S1

#### Figure 5- supplement 1

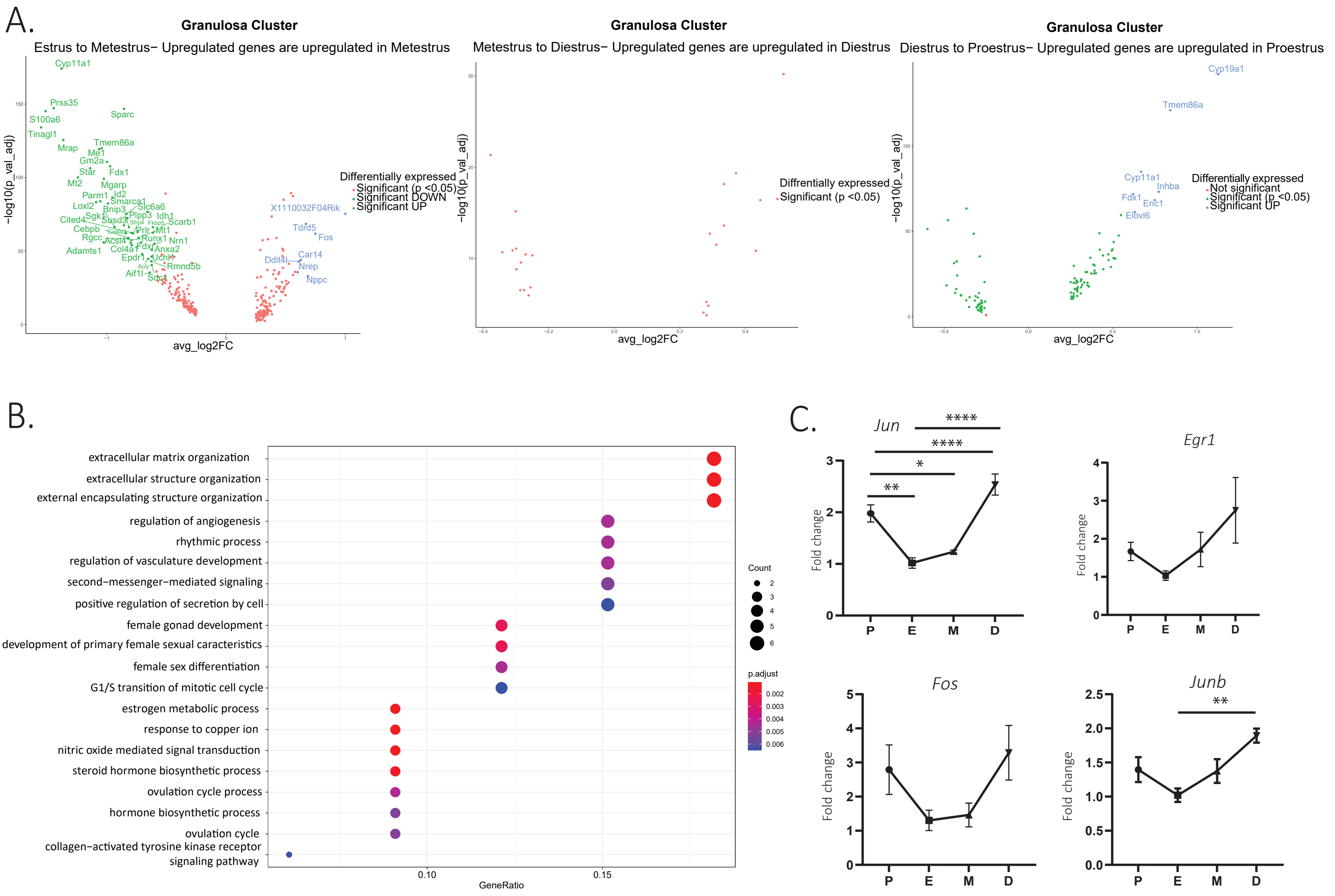
