## Supplementary file 1 for "A single cell atlas of the cycling murine ovary"

| **Supplementary File 1- List of primers used for qPCR experiments** | | |  |
| --- | --- | --- | --- |
| Gene | Forward | Reverse | NCBI |
| Cyp19a1 | TCCACACTGTTGTGGGTGAC | AGGGAAGTACTCGAGCCTGT | NM_007810.4 |
| Egr1 | GGAGAGGCAGGAAAGACATAAA | GCTCTGAGATCTTCCATCTGAC | NM_007913.5 |
| Fos | AATGGTGAAGACCGTGTCAG | GGATTCTCCGTTTCTCTTCCTC | NM_010234.3 |
| Gapdh | GGGGCTCTCTGCTCCTCCCTG | CCAGGCGTCCGATACGGCCA | NM_001289726.1 |
| Inhba | GCTGCTCAAGTGCCAATACC | GTTTTGTCAGCCGGCTCTTG | NM_008380.2 |
| Jun | CCAGACTGTACACCAGAAGATG | CAACCAAAGTGTCTGCTTTCC | NM_010591.2 |
| Junb | TCACGACGACTCTTACGCAG | CCTTGAGACCCCGATAGGGA | NM_008416.3 |
| Lhcgr | GAGTGATTCCCTGGAAAGGATAG | AGCACCGGGTTCAATGTATAG | NM_001364898.1 |
| Nppc | GCGGTCTGGGATGTTAGTG | CATTGCGTTGGAGGTGTTTC | NM_010933.5 |
| Pgr | GGTGCGAGGATTCTGACATT | TATGGCTCCCTAGGTTCTCTAC | NM_008829.2 |
| Prss35 | CAGTAAGCATCTCAGAGGGCA | GGGGCTGGCAAGATGGATAG | NM_178738.3 |
| Rgcc | ACTCCTCGGAAAGCCAAATTA | CCAAACTCCTTGCTTCACATAC | [NM_025427.2](https://www.ncbi.nlm.nih.gov/nuccore/NM_025427.2) |
| Sgk1 | AGCCATCCTGAAGAAGAAAGAG | GGTCTGGAATGAGAAGTGAAGG | NM_001161845.2 |
| Star | GCTGTGAAGGCTAAGGGATAAG | GTGACATTTGGAGCTGGTAAGA | NM_011485.5 |
| Tinagl1 | GAATCCCACCTAGGAGACAAAG | GTCTATACTCCCTGCAGTTTCC | [NM_001168333.1](https://www.ncbi.nlm.nih.gov/nuccore/NM_001168333.1) |
| Trib2 | TTGGAAACTGCAGGGAGTATAG | ATGAGAGAGAGGGTGGGTAG | NM_001168333.1 |
