## Supplementary File 2 for "A single cell atlas of the cycling murine ovary"

**Supplementary File 2 - Top 10 markers expressed in each ovary cluster**

| **Granulosa** | avg_log2FC | **Mesenchyme** | avg_log2FC | **Endothelium** | avg_log2FC |
| --- | --- | --- | --- | --- | --- |
| Nr5a2 (88) | 2.37 | Col1a2 (89) | 2.80 | Kdr (90) | 3.21 |
| Bex4 (4) | 2.08 | Col3a1 (91) | 2.68 | Mmrn2 (92) | 3.15 |
| Lect1 (93) | 1.95 | Col1a1 (4) | 2.64 | Cdh5 (4) | 3.02 |
| Fst (94) | 1.90 | Ogn (68,73) | 2.40 | Egfl7(95) | 2.93 |
| Hsd17b1 (96) | 1.89 | Bgn (97) | 2.37 | Pecam1 (98) | 2.91 |
| Slc18a2 (8,99) | 1.87 | Tcf21 (100) | 2.35 | Cldn5 (101) | 2.87 |
| Inha (8) | 1.82 | Dcn (29) | 2.34 | Esam (102) | 2.84 |
| Serpine2 (103) | 1.76 | Pdgfra (104) | 2.30 | Flt1 (105) | 2.83 |
| Ivns1abp (106) | 1.65 | Lum (73) | 2.18 | Cd93 (92,107) | 2.79 |
| Fam13a | 1.59 | Mgp (108) | 2.17 | Ctla2a (109) | 2.77 |

| **Immune** | avg_log2FC | **Oocyte** | avg_log2FC | **Epithelilum** | avg_log2FC |
| --- | --- | --- | --- | --- | --- |
| Lyz2 (110) | 3.62 | Gm15698 | 4.06 | Upk1b (111) | 3.23 |
| Laptm5 (112) | 3.54 | Gdf9 (113) | 3.91 | Upk3b (114) | 3.17 |
| H2Aa (115) | 3.48 | H1foo (116) | 3.78 | Lgals7 (117) | 3.09 |
| Cd74 (9) | 3.47 | Padi6 (118) | 3.78 | Aldh1a2 (119) | 3.04 |
| Fcer1g (120) | 3.45 | Ooep (121) | 3.75 | Mt2 (122,123) | 2.95 |
| Ctss (9) | 3.37 | Oosp1 (124) | 3.72 | Ildr2 (125) | 2.90 |
| H2.Eb1 (126) | 3.33 | Rfpl4 (127) | 3.62 | Krt18 (75) | 2.82 |
| C1qb (128) | 3.32 | Tcl1 (124) | 3.56 | Gpm6a | 2.76 |
| C1qc (128) | 3.23 | Khdc1b (129) | 3.54 | Plxna4 (130) | 2.75 |
| Cd52 | 3.17 | Nlrp14 (131) | 3.53 | Krt7(75) | 2.67 |
