## Supplementary File 3 for "A single cell atlas of the cycling murine ovary"

**Table S3 - Top 10 markers from each mesenchyme subclusters**

| **Steroidogenic stroma** | avg_log2FC | **Fibroblast-like stroma** | avg_log2FC | **Early theca** | avg_log2FC |
| --- | --- | --- | --- | --- | --- |
| Tenm4 | 1.43 | Enpp2 (132) | 2.09 | Enpep (133) | 1.72 |
| Htra3 (134) | 1.26 | Fzd1 (23) | 1.97 | Adcy7 (135) | 1.64 |
| Ogn | 1.16 | Gatm (23) | 1.95 | Thbd (136) | 1.52 |
| Tcf21 (137) | 1.07 | Igfbp4 (138) | 1.62 | Mest (9) | 1.37 |
| Itih5 | 1.05 | Tnfrsf19 (139) | 1.62 | Stmn1 | 1.35 |
| Mfap4 | 1.26 | Plat | 1.45 | Mycn (2) | 1.31 |
| Rarres2 (140) | 1.03 | Cadm4 (141) | 1.44 | Hhip (68) | 1.31 |
| Pmepa1 (142) | 1.04 | Lpl (143) | 1.50 | Lamc3 | 1.26 |
| Kcnk2 (144) | 1.05 | Tnc (145) | 1.44 | Hmgb2 | 1.30 |
| Cxcl12 (146,147) | 1.14 | Tgfbi | 1.36 | Top2a | 1.28 |

| **Steroidogenic theca** | avg_log2FC | **Smooth muscle** | avg_log2FC | **Pericyte** | avg_log2FC |
| --- | --- | --- | --- | --- | --- |
| Mgarp (29) | 2.90 | Myh11 (9) | 2.98 | Rgs5 (148) | 3.76 |
| Aldh1a1 (29,68,68) | 2.50 | Actg2 (149) | 2.82 | Notch3 (150) | 3.00 |
| Cyp17a1 (2) | 2.48 | Cnn1 (151) | 2.68 | Ndufa4l2 | 2.79 |
| Cyp11a1 (9) | 2.44 | Tagln (9,152) | 2.46 | Rgs4 | 2.65 |
| Fdx1 (9,153) | 2.36 | Ednrb (154) | 2.26 | Gm13889 | 2.63 |
| Hao2 | 2.33 | Lmod1 (155) | 2.22 | Ebf1 | 2.50 |
| Folr1 (156) | 2.21 | Fhdc1 | 2.05 | Steap4 | 2.41 |
| Acsbg1 | 2.12 | Fhl1 (157) | 2.03 | Serpini1 | 2.40 |
| Serpina5 (158) | 2.07 | Tpm2 (159) | 2.01 | Tinagl1 | 2.40 |
| Dnajc15 | 2.03 | Acta2 (2,9) | 2.01 | Gja4 | 2.38 |
