## Supplementary File 4 for "A single cell atlas of the cycling murine ovary"

**Supplementary File 4 - Top 10 markers from each granulosa subclusters**

| **Preantral-Cumulus** | avg_log2FC | **Antral-Mural** | avg_log2FC | **Atretic** | avg_log2FC | **Mitotic** | avg_log2FC |
| --- | --- | --- | --- | --- | --- | --- | --- |
| Gatm (9) | 2.13 | Inhba (99,160) | 2.25 | Ghr | 1.85 | Top2a (161) | 2.30 |
| Kctd14 (8) | 2.05 | Nppc(80,162,163) | 2.01 | Pik3ip1 (164) | 1.80 | Ube2c (165) | 2.11 |
| Igfbp5 (8) | 1.87 | Mro (61,166) | 2.00 | Cald1 (167) | 1.78 | Racgap1 (39,106) | 1.97 |
| Col18a1 | 1.86 | Nap1l5 (168) | 1.70 | Itih5 | 1.63 | Birc5 | 1.96 |
| Pcsk6 (169) | 1.85 | Hsd17b1 (9) | 1.59 | Cfh | 1.61 | Ccna2 (170,171) | 1.95 |
| Slc18a2 (31) | 1.84 | Slc26a7 (93) | 1.56 | Asb4 | 1.56 | Ccnb2 | 1.86 |
| Fndc5 (172) | 1.79 | Grem1 (99) | 1.54 | Ctgf | 1.51 | Hmgb2 (173) | 1.85 |
| Wt1 (2,9,174–176) | 1.78 | X1110032F04Rik | 1.53 | Ntn4 (167) | 1.51 | Cdca8 (177) | 1.81 |
| Tmem184a (8) | 1.77 | Cyp19a1 (178) | 1.52 | Cdc42ep3 | 1.44 | Cdk1 (59,179) | 1.80 |
| X1190002N15Rik | 1.74 | Tom1l1 | 1.39 | Fhl1 | 1.40 | Prc1 | 1.75 |

| **Luteinizing mural** | avg_log2FC | **Active CL** | avg_log2FC | **Mitotic-Antral** | avg_log2FC | **Regressing CL** | avg_log2FC |
| --- | --- | --- | --- | --- | --- | --- | --- |
| Tinagl1 (82) | 2.90 | Neat1 (33) | 2.13 | Ccnb2 (180,181) | 1.94 | Ptgfr (167,182) | 2.74 |
| Adamts1 (61,62) | 2.50 | Col3a1 (99) | 2.09 | Ube2c (165) | 1.83 | Efhd1 (183) | 2.50 |
| Mrap (167) | 2.45 | Igfbp7 (184) | 2.03 | Cenpa | 1.75 | S100a6 (185,186) | 2.47 |
| Prss35 (60) | 2.40 | Sfrp4 (187,188) | 1.82 | Top2a (161) | 1.74 | Tnc (61,187) | 2.42 |
| Mt2 (59,96,189) | 2.30 | Cyp11a1 (58) | 1.70 | Racgap1 (39,106) | 1.73 | Lipg (190) | 2.37 |
| Parm1 (191) | 2.26 | Plin4 | 1.71 | Birc5 | 1.72 | Sfrp4 | 2.28 |
| Loxl2 (192) | 2.25 | Onecut2 | 1.71 | Inhbb (9,99) | 1.62 | Lgmn | 2.27 |
| S100a6 (186) | 2.22 | Col1a2 (89) | 1.96 | Ccna2 (170,171) | 1.61 | Star | 2.25 |
| Cemip | 2.20 | Col1a1 (4) | 1.84 | Nap1l5 (168) | 1.60 | Akr1c18 (96,193) | 2.24 |
| Cdkn1a | 2.14 | Gm42669 | 1.75 | Cdca8 (177) | 1.59 | Sgk1 | 2.21 |
