## Supplementary File 5 for "A single cell atlas of the cycling murine ovary"

**Table S5 – Secreted markers expressed in granulosa cells varying with the estrous cycle**

| Aebp1 | Adamts1 | Col4a2 | Inhba | Prss35 |
| --- | --- | --- | --- | --- |
| Bmper | Anxa2 | Col6a1 | Inhbb | Sparc |
| Htra1 | Bmp2 | Cyr61 | Igfbp7 | Sfrp4 |
| Htra3 | Ctsb | Crispld2 | Ltbp1 | Serpine2 |
| Ihh | Ctsd | Epdr1 | Lgals1 | Sbsn |
| Npc2 | Cemip | Fbln1 | Lect1 | Sdc1 |
| Smoc2 | Clu | Fst | Loxl2 | Tnc |
| Sparcl1 | Col1a2 | Gsn | Nppc | Timp1 |
| St3gal1 | Col3a1 | Grem1 | Pcolce | Tinagl1 |
| Ybx1 | Col4a1 | Inha | Psap | Vegfa |
